# Time-structured communication through cross-frequency bursts

**DOI:** 10.64898/2026.04.30.721821

**Authors:** Jung Young Kim, Sang Wan Lee, Jee Hyun Choi, Demian Battaglia

## Abstract

Adaptive brain function requires communication that is selective in both space and time, yet the circuit mechanisms that transiently favor specific routes of information transfer remain unclear. Here we show that, when coupled, spiking populations with distinct inhibitory timescales can reprogram one another’s intrinsic burst preferences, generating irregular multi-frequency bursts that self-organize into Multi-Frequency Oscillatory Patterns (MFOPs). State-resolved transfer entropy revealed that different MFOPs were associated with distinct Information Routing Patterns (IRPs), defining transient windows of enhanced directed transfer with specific delays and directions. At the microscopic level, the same oscillatory states selectively increased or decreased delayed spike transmission across monosynaptic pathways, yielding transmission barcodes that significantly resembled the corresponding IRPs. This repertoire was markedly reduced when inhibitory diversity was removed. These results identify coordinated multi-frequency bursting as a mechanism by which recurrent circuits can dynamically filter inputs according to source and latency.

## Introduction

Adaptive brain function requires flexible communication across distributed circuits, with inputs dynamically weighted by source and timing so that the right signals reach the right targets at the right moment. The representation and processing of complex environments, behaviors and contexts therefore require spatiotemporal control of neuronal activity—not rate variation alone [1–3]. Yet the mechanisms that gate information exchange across space and time remain incompletely understood.

Proposed mechanisms include precise inhibitory timing within local circuits [4], dendritic nonlinearities shaping temporal integration [5], temporal filtering by short-term synaptic adaptation [6], flexible interactions across multiple interareal pathways [7], shifts in brain state [8], and modulation by propagating activity fronts [9]. Among these hypotheses, cross-frequency oscillations are particularly appealing as a potential scaffold for organizing when and where information is encoded and transmitted [10]. Spatial and non-spatial information can be encoded in distinct fast cycles nested within a slower rhythm [11, 12], with different combinations of nested frequencies associated with different types of information [13].

Beyond encoding, oscillations may also create temporal windows for dynamic information exchange across populations through flexible phase alignment [14, 15]. This communication-through-coherence framework has been extended to coexisting oscillations at multiple frequencies, suggesting that different bands may mediate multiplexed transfer from distinct sources [16, 17] or in different directions [18, 19], thereby supporting hierarchical predictive computations [20]. Frequency bands may also interact to gate one another, favoring alternative directed routes at different times [21, 22].

However, this framework faces an important tension. Neural rhythms in vivo are typically not sustained metronomes, but transient, stochastic and burst-like events [23, 24]. Their amplitudes, durations and instantaneous frequencies often drift substantially from cycle to cycle [25–27]. This irregularity raises a deeper question than whether oscillatory communication merely remains possible under noisy conditions. If cross-frequency structure emerges only in brief and variable bursts, can it still organize communication sequentially in time, creating reproducible windows in which different routes, targets or directions of influence are transiently favored? Or do theories of oscillatory multiplexing rely too heavily on idealized, stationary band-limited rhythms?

Here, we model neuronal populations that spontaneously generate irregular, transient bursts. Assigning distinct synaptic time constants to different interneuron classes reproduces the diversity of intrinsic bursting frequencies associated with inhibitory cell-type heterogeneity [28–30]. In isolation, individual modules oscillate at their own characteristic frequencies. Yet once coupled through excitatory and inhibitory connections, these intrinsic rhythms reorganize collectively: circuits tuned to fast oscillations can also generate slow bursting, and vice versa. This self-organized frequency reprogramming, beyond structural hardwiring, gives rise to a rich landscape of transient Multi-Frequency Oscillatory Patterns (MFOPs).

Extending previous information-theoretic analyses of gamma bursts [15], we show that coordinated multi-frequency bursts transiently organize information routing. These events form Information Routing Patterns (IRPs): sequences of enhanced transfer structured in both space and time, akin to temporal motifs in communication networks [31, 32]. At the spike level, bursts create alternating windows of increased and decreased transmission probability, paced by the slowest frequency component.

Our results challenge the idea that each frequency band maps onto a fixed communication channel. Even brief, stochastic bursts can reliably gate transmission through multi-frequency coordination, providing fine-grained temporal flexibility—like rhythmic accents emphasizing different words in a verse. Crucially, this flexibility depends on inhibitory diversity, and is markedly reduced when such diversity is removed.

Thus, cross-frequency interactions emerge as a general organizing principle, not merely for multiplexed routing, but for the spatiotemporal sculpting of relevance masks, conceptually parallel to positional encoding and attentional masking in artificial neural networks [33, 34].

## Results

### Distinct inhibitory timescales generate transient bursts at different intrinsic frequencies

Frequency diversity across brain circuits is often attributed, at least in part, to differences in inhibitory microcircuit dynamics and in the interneuron classes that dominate them [29, 30]. Motivated by this idea, we began by constructing two local spiking-network populations that differ only in the time constants of their inhibitory synapses (Methods), thereby hardwiring different intrinsic oscillatory tendencies while keeping the architecture otherwise identical. We refer to these populations as `Pop`_*F*_ and `Pop`_*S*_.

When simulated in isolation, both populations operated in a fluctuation-driven regime in which collective oscillations were not sustained, but emerged intermittently as brief synchronization events (Fig. 1a,b). Raster plots revealed sparse spiking punctuated by transient episodes of coordinated firing, and these events were mirrored in the population-averaged membrane potential, which displayed short-lived oscillatory waveforms rather than continuous rhythmic activity. Despite this common bursty regime, the two populations expressed clearly different intrinsic frequencies: bursts in `Pop`_*F*_ were centered at higher frequencies (Fig. 1a), whereas bursts in `Pop`_*S*_ remained in a lower range (Fig. 1(b)). Thus, simply changing inhibitory synaptic timescales was sufficient to generate two populations with distinct native oscillatory preferences while preserving the transient character of their activity.

**Figure 1.**
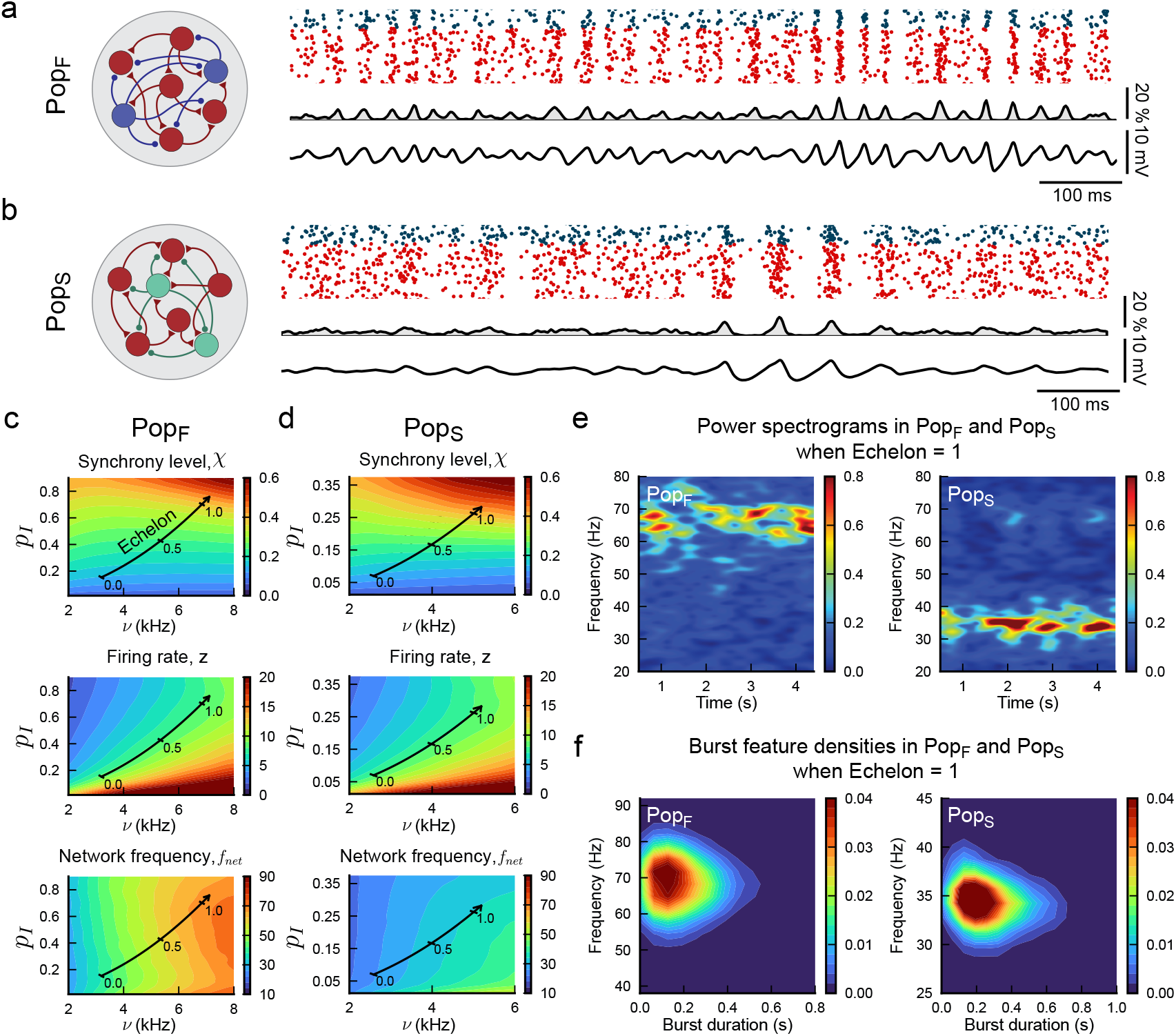
Isolated populations with distinct inhibitory timescales generate transient bursts at different intrinsic frequencies. **a**. Schematic of the Pop_*F*_ (left) and representative simulated activity in the transient-bursting regime (right). Raster plot shows excitatory (red) and inhibitory (blue) neurons (top), multiunit activity (middle), and population-averaged membrane potential used as an LFP-like signal (bottom). Activity is sparse and irregular, but intermittently organizes into short-lived synchronized events (at a 30–40 Hz frequency). **b**. Same as in a for the `Pop`_*S*_. Bursting remains transient, but occurs at a lower intrinsic frequency (10–15 Hz) than in the fast Population. **c**, Activity statistics of the `Pop`_*F*_ across inhibitory connection probability and external input rate. Top, synchrony level *χ*. Middle, firing rate *z*. Bottom, network frequency *f*_*net*_. The black curve indicates the iso-firing-rate trajectory used to define the echelon coordinate, along which synchrony increases while firing rate remains approximately constant. **d**. Same as in c for the `Pop`_*S*_. The two populations span similar ranges of synchrony and firing rate, but differ in intrinsic oscillation frequency. **e**. Representative spectrograms of the `Pop`_*F*_ (left) and `Pop`_*S*_ (right) at echelon = 1. In both cases, oscillatory activity is organized into brief bursts rather than sustained rhythms, with lower frequencies in the `Pop`_*S*_ and higher frequencies in the `Pop`_*F*_. **f**. Burst-feature densities at echelon = 1 for the `Pop`_*F*_ (left) and `Pop`_*S*_ (right), showing the joint distribution of burst duration and frequency. Bursts are short-lived in both populations, but concentrated at distinct frequency ranges.

To characterize these regimes systematically, we mapped the activity of each isolated population across inhibitory connection probability and external drive, and quantified three summary statistics: synchrony level (*χ*), firing rate (*z*), and network frequency (*f*_*net*_) (Methods). In both populations, increasing synchrony required coordinated changes in these parameters, whereas firing rate and synchrony were not independently controlled a priori (Fig. 1c,d). We therefore sought a parametrization that would allow us to vary the strength of collective coordination without introducing a confound from mean firing rate.

For this purpose, we identified, in each population separately, an iso-firing-rate trajectory through parameter space along which synchrony increased smoothly while firing rate remained approximately constant (Fig. 1c,d; Fig. S1a). We denote position along this trajectory by an effective coordinate, the echelon, ranging from 0 to 1. The echelon therefore provides a one-dimensional control parameter that increases collective synchrony through a multiparameter change, while minimizing differences in average spiking output between conditions.

Along these iso-rate trajectories, both populations remained in a partially synchronized, burst-dominated regime even at the highest echelon values. Higher echelons increased synchrony and sharpened the oscillatory signature, but did not convert the dynamics into stable coherent rhythms (Fig. S1a,b). Instead, spectrograms and burst-feature densities showed that activity was still organized into brief oscillatory episodes, typically lasting less than 0.5 s, with distinct frequency ranges preserved across populations: approximately 60–80 Hz in `Pop`_*F*_ and 30–40 Hz in `Pop`_*S*_ at echelon = 1 (Fig. 1e,f; Fig. S1b). Decreasing echelon reduced burst prominence and shifted frequencies downward in both populations, again without abolishing the transient nature of the oscillations.

These analyses establish a controlled baseline for the rest of the study. By construction, `Pop`_*F*_ and `Pop`_*S*_ differ in intrinsic burst frequency, yet can be matched in average firing rate and probed at comparable levels of synchrony. This separation of timescale from firing-rate effects provides the starting point for asking how interactions between populations with different native bursting tendencies reshape oscillatory coordination and, ultimately, information routing and input integration.

### Coupling reprograms intrinsic burst preferences into distinct multi-frequency regimes

Having established two isolated populations with distinct intrinsic burst frequencies, we next asked how these native oscillatory tendencies are altered once the populations interact. This question is central because frequency differences across cortical circuits are often attributed to local microcircuit composition, in particular to the interneuron classes that shape local resonance [29, 30]. Yet putative oscillation generators are never perfectly segregated in real circuits. Even when distinct rhythms are preferentially associated with different layers, interneuron types or subnetworks, these components remain embedded in the same tissue and remain in continuous reciprocal interaction through excitation and inhibition. Their oscillatory identities should therefore not be regarded as fixed local signatures, but as properties reshaped by network interaction itself.

To capture this idea, we coupled `Pop`_*F*_ and `Pop`_*S*_ through both excitatory and inhibitory projections (Fig. 2a). This choice was motivated in part by an interlaminar interpretation of the two modules: if the populations are viewed as local circuits embedded in different cortical layers, their reciprocal interactions need not be purely excitatory, but also be inhibitory, because of diffuse recurrent inhibition. Three structural parameters controlled this inter-population architecture: *α* scaled excitatory coupling, *β* scaled inhibitory coupling, and *ω* set their directional asymmetry (Methods). As in the previous section, the echelon controlled within-population synchrony while maintaining approximately constant firing rate. Together, these parameters define a family of coupled circuits in which populations with different hardwired intrinsic timescales interact with variable strength and directional bias.

**Figure 2.**
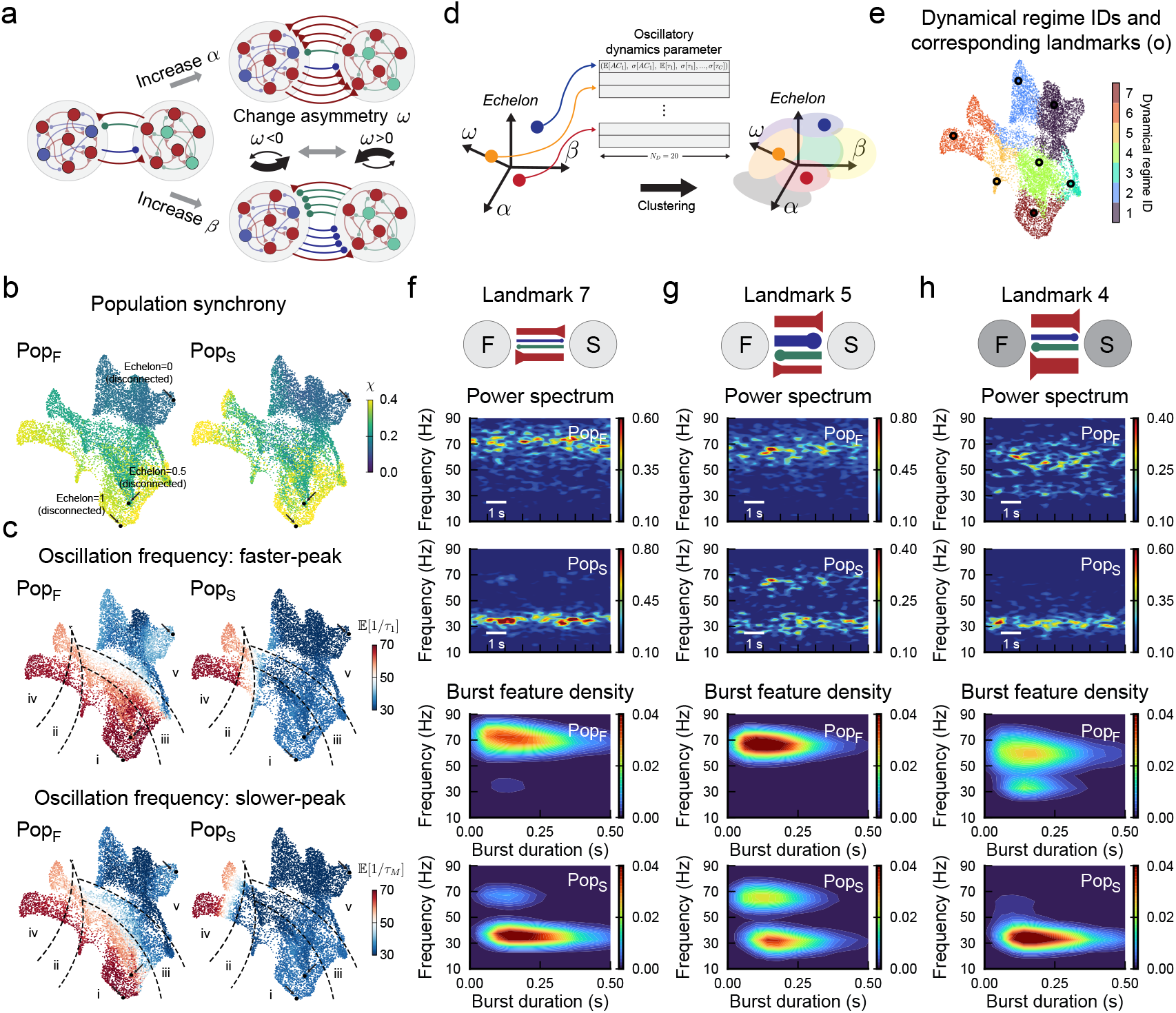
Coupling reprograms intrinsic burst preferences into distinct multi-frequency dynamical regimes. **a**. Schematic of the coupled two-population circuit. The parameter *α* scales excitatory inter-population coupling, *β* scales inhibitory inter-population coupling, and *ω* controls directional asymmetry (Methods). Echelon modulates within-population synchrony while maintaining approximately constant firing rate. **b**. UMAP embedding of coupled-circuit dynamics based on oscillation-related dynamical features extracted from both populations. Each point corresponds to one structural parameter set. Point color indicates synchrony level for the fast (left) and slow (right) populations. Black markers indicate the disconnected reference cases at echelon = 0, 0.5 and 1. **c**. Same embedding colored by the primary and secondary bursting frequencies expressed in each population. These frequencies were inferred from the multi-peak structure of the population autocorrelograms. When only one bursting component was present, primary and secondary frequencies coincided. Dashed contours indicate broad sectors with qualitatively distinct oscillatory behavior. **d**. Schematic of the clustering procedure used to group circuit configurations with similar oscillatory dynamics based on the extracted feature set. **e**. Dynamical regime identity mapped onto the UMAP embedding. Seven regimes were identified by unsupervised clustering. Black markers indicate landmark configurations selected as representative examples of each regime. **f–h**. Representative landmarks illustrating distinct classes of coupled oscillatory dynamics. Top, schematic of the corresponding coupling architecture; red arrows indicate excitatory projections, blue inhibitory projections from `Pop`_*F*_ to `Pop`_*S*_, and green inhibitory projections from `Pop`_*S*_ to `Pop`_*F*_. Arrow width scales with projection ratio, and node shading indicates echelon. Middle, representative spectrograms for the `Pop`_*F*_ and `Pop`_*S*_. Bottom, joint burst-duration/frequency densities for both populations. **f**, Landmark 7 largely preserves the native frequency contrast between populations. g, Landmark 5 shows a secondary fast bursting component in the `Pop`_*S*_. **h**, Landmark 4 shows a secondary slow bursting component in the `Pop`_*F*_ .

For each circuit configuration, we simulated spontaneous activity and extracted a set of oscillation-related dynamical features from the two populations and from their cross-correlation structure (Methods; Fig. S2). We then used these features to construct a low-dimensional embedding of the dynamical landscape (Fig. 2b; full overlays in Fig. S3). Each point in this space corresponds to one coupled circuit configuration, with nearby points representing similar burst dynamics. As in the disconnected case, echelon was a major organizing variable: increasing synchrony displaced the dynamics along a dominant axis of the embedding. Yet synchrony alone did not determine the oscillatory regime. At comparable synchrony levels, changing the pattern of inter-population coupling could move the system to distant regions of the map, especially for the slow population, indicating that coupling reshapes burst organization in ways that cannot be reduced to a simple modulation of coherence.

We next examined how the bursting frequencies expressed by each population varied across this landscape. To do so, we tracked the primary and secondary bursting frequencies inferred from the multi-peak structure of the population autocorrelograms (Fig. 2c; Fig. S2). The primary frequency corresponds to the dominant bursting timescale of a population, whereas the secondary frequency captures an additional oscillatory component when present. In some zones of the oscillatory dynamics map, the coupled system largely preserved its native organization: `Pop`_*F*_ remained dominated by a fast primary bursting frequency, whereas `Pop`_*S*_ retained a slower one (e.g., zone *i* in the low dimensional map of Fig. 2b). In other regions, one population acquired a secondary bursting frequency near the preferred timescale of the other while preserving its own primary mode (e.g., region *ii* and *iii*). Thus, `Pop`_*S*_ could express a secondary fast component in addition to its native slow bursting, or conversely `Pop`_*F*_ could acquire a secondary slow component. In still other regions, the primary bursting frequencies of the two populations converged, such that both became dominated by a common slow or a common fast mode. The coupled system therefore does not merely superimpose two pre-existing rhythms. Rather, it generates a repertoire of multi-frequency states in which intrinsic burst preferences may be preserved, hybridized or effectively reassigned. Note that the color coding in Fig. 2c represents the primary and secondary bursting frequencies inferred from averaged autocorrelograms. In some regions (e.g. region *i*), the similar color of primary and secondary frequencies indicates a strong dominance of one frequency, but does not rule out rarer bursts at other frequencies that are too infrequent to produce clear autocorrelogram peaks. Frequency variability and reprogramming may therefore be richer than suggested by the simplified representation in Fig. 2b.

Although this landscape was diverse, it was not unstructured. We therefore asked whether the observed behaviors could be partitioned into a limited number of dynamical regimes. To this end, we applied unsupervised clustering to the oscillation-related feature set (Fig. 2d; diagnostics in Fig. S4). This analysis identified seven robust regimes, which occupied compact and partially separated regions of the embedding (Fig. 2e). From each regime, we selected one representative circuit configuration, referred to as a landmark, for closer inspection (see Methods).

These landmarks make the logic of the coupled system especially clear. Landmark 7 corresponds to a regime in which the original frequency contrast between the populations is largely preserved: `Pop`_*F*_ exhibits a fast primary bursting frequency around 70 Hz, whereas `Pop`_*S*_ remains centered near 30–40 Hz, with bursts in both populations still transient and typically shorter than 0.5 s (Fig. 2f). Landmark 5 illustrates a different configuration, in which `Pop`_*S*_ acquires a clear secondary fast bursting component while retaining a slow primary one, whereas `Pop`_*F*_ remains primarily dominated by the fast mode (Fig. 2g). Landmark 4 shows the complementary situation: `Pop`_*F*_ expresses both slow and fast bursting components, with a secondary slow component superimposed on its fast primary mode, whereas `Pop`_*S*_ remains mainly in the slower range (Fig. 2h). Additional landmarks further illustrate regimes in which both populations converge toward similar primary frequencies or jointly express mixed multi-frequency bursting (Fig. S5).

Together, these results show that coupling creates a structured repertoire of transient multi-frequency regimes that cannot be inferred from the disconnected populations alone. Hardwired local timescales still matter, but they no longer rigidly determine the oscillatory identity of each population once interactions are engaged. Instead, inter-population coupling can preserve, mix or override intrinsic burst preferences, making oscillatory structure an emergent property of the circuit as a whole. This repertoire of coupled burst states, with effectively reprogrammed intrinsic properties, provides the dynamical substrate on which time-structured communication can be selectively routed.

### Bursting self-organizes into a structured repertoire of cross-population co-bursting patterns

The coupled regimes described above are not defined only by the frequencies each population can express, but also by how burst events become coordinated across populations. A crucial question is therefore whether the fast and slow bursting components identified within a given regime arise largely independently, or instead tend to occur in temporally matched configurations. This distinction matters because routing can only depend on oscillatory state if the relevant burst events form coordinated circuit-level patterns rather than isolated local fluctuations.

To address this point, we quantified burst co-occurrence across frequencies and populations. For each simulation, we first identified transient burst events by thresholding the spectrogram separately at each frequency (Methods), yielding a binary time-frequency mask for each population (Fig. 3a,b). We then measured, for every pair of frequencies across the two populations, the probability that burst events were simultaneously present. An example landmark (here landmark 7) is shown in Fig. 3, to illustrate the procedure and the type of structure that emerges. In this example, the co-occurrence map displays clear peaks rather than a diffuse background (Fig. 3c). As expected, strong time-aligned co-bursting occurs between matching slow components across populations. However, significant peaks also link different frequencies, showing that burst coordination is not restricted to same-frequency synchrony. Instead, slow bursts in one population can be preferentially matched to fast bursts in the other, and vice versa. Thus, the relevant circuit states are not defined by isolated bursting within each population, but by recurrent patterns of coordinated multi-frequency activation.

**Figure 3.**
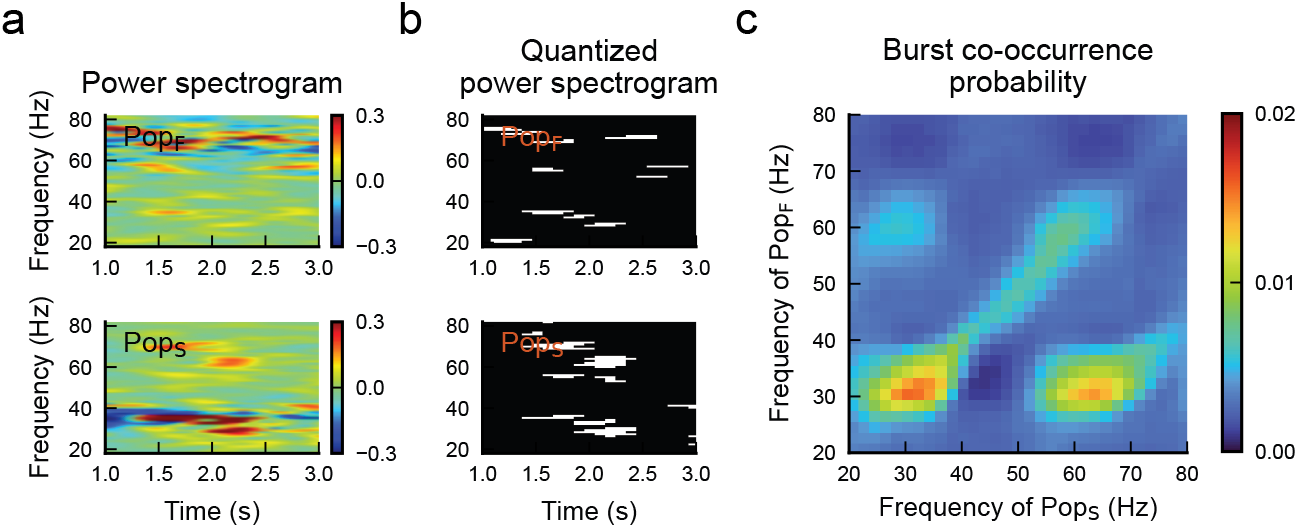
Multi-frequency bursts form coordinated co-bursting patterns across populations. **a**. Representative power spectrograms from the `Pop`_*F*_ (top) and `Pop`_*S*_ (bottom) for one example landmark, shown here only to illustrate the analysis procedure. Distinct burst components are visible in both populations over time. **b**. Quantized spectrograms obtained by thresholding oscillatory power separately at each frequency (Methods), yielding binary time-frequency masks that identify transient burst events in the `Pop`_*F*_ and `Pop`_*S*_. **c**. Burst co-occurrence probability between frequencies in the two populations, computed from the overlap of the quantized spectrograms in **b**. Peaks indicate frequency pairs whose bursts co-occur more often than expected from isolated local bursting. In the example shown, co-bursting is enhanced not only between matching slow components, but also across different frequencies. See Fig. S6 for the corresponding maps across all landmarks and significance assessment.

This phenomenon was not specific to the example shown. Across landmarks, co-bursting maps consistently revealed structured peaks within populations and between populations, many of them significantly above surrogate expectations (Fig. S6). Different landmarks favored different combinations, but the general picture was the same: multi-frequency bursting was organized into repeated joint configurations rather than arising from independent local events. In other words, once the two populations are coupled, oscillatory complexity is no longer purely local, but becomes jointly coordinated, as a “free-lunch” from self-organization. What matters is not only which frequencies each population can generate, but which combinations of bursts tend to recur together across the circuit.

These frequently reproduced co-bursting configurations provide a natural unit for the next stage of the analysis. We therefore define a *Multi-Frequency Oscillatory Pattern* (MFOP) as the combination of a given landmark and a specific coordinated burst configuration across populations and frequencies (Fig. 4a). MFOPs thus capture characteristic oscillatory transients that emerge within each dynamical regime: they are not merely labels of local oscillatory content, but characteristic signatures of coordinated multi-frequency bursting. This definition allows us to move from the space of dynamical regimes to the finer-grained space of transient oscillatory states, and to ask whether different co-bursting patterns support different routing configurations.

**Figure 4.**
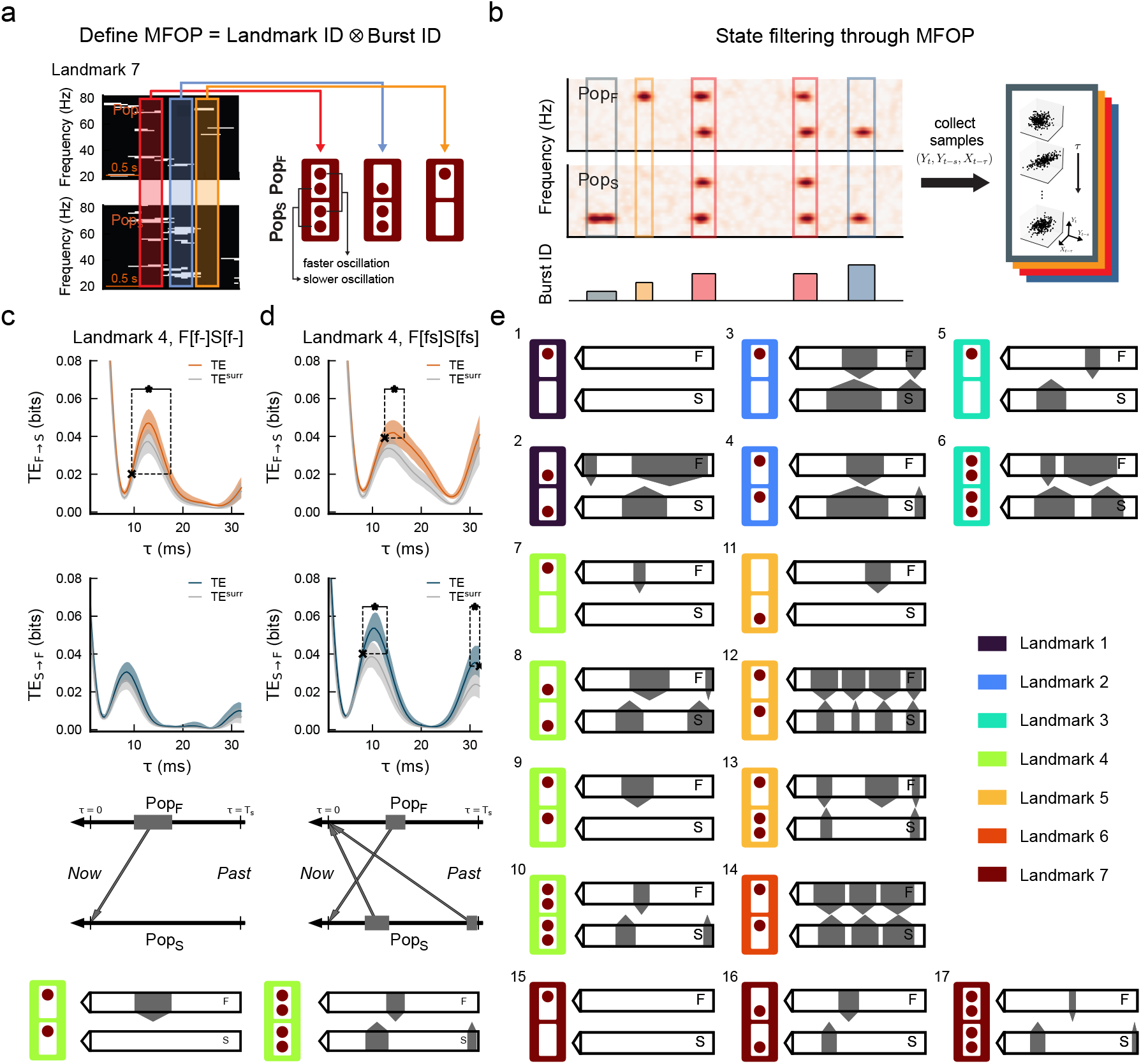
State filtering by coordinated multi-frequency bursts reveals Information Routing Patterns. **a**. Schematic definition of a Multi-Frequency Oscillatory Pattern (MFOP). An MFOP is defined as the combination of a landmark identity and a coordinated burst mask across the `Pop`_*F*_ and `Pop`_*S*_. Example pictograms are shown for landmark **b**. State filtering through MFOPs. Time points at which a given MFOP is active are collected across simulations to assemble MFOP-specific samples of population activity. Directed functional connectivity is then estimated within each state rather than in short sliding windows. **c**. Example lagged TE profiles and Information Routing Pattern (IRP) construction for one MFOP from landmark 4. Top, TE from `Pop`_*F*_ to `Pop`_*F*_ as a function of transmission delay *τ* ; middle, TE from `Pop`_*S*_ to `Pop`_*F*_ ; gray curves indicate surrogate TE. The shaded region indicates the 5th–95th percentile range. Asterisks mark TE peaks exceeding surrogate expectations, and crosses indicate the return to surrogate level used to delimit the corresponding boosted-transfer interval. Bottom, schematic summary of the resulting transfer window and directionality, yielding the IRP associated with the MFOP. **d**. Same analysis as in **c** for a different MFOP within the same landmark. Although the two MFOPs belong to the same structural-dynamical regime, they give rise to distinct lagged transfer profiles and therefore distinct IRPs. **e**. Diversity of IRPs identified across MFOPs. Colored frames indicate landmark identity. Each diagram summarizes the temporal windows and directions of boosted transfer associated with one MFOP.

### State filtering by coordinated multi-frequency bursts reveals Information Routing Patterns

Having identified MFOPs as recurrent states of coordinated multi-frequency bursting, we next asked whether different MFOPs support different patterns of inter-population communication. To address this question, we quantified directed functional connectivity using lagged Transfer Entropy (TE) between the multi-unit activities of the two populations [35]. TE measures how much information the past activity of one population provides about the present activity of the other beyond the target’s own past, and thus yields a model-free estimate of directed information transfer at delay *τ* (see Methods).

Our analysis is state-resolved rather than simply time-resolved. Instead of estimating TE in short sliding windows and then following its fluctuations over time, we first identified all time points at which a given MFOP was active, pooled those samples across long simulations, and then estimated TE conditional on that state (Fig. 4a,b). This extends the state-filtering strategy introduced by Palmigiano et al. [15] to the present multi-frequency setting, in which states are defined not merely by the presence or absence of a burst, but by coordinated burst masks across frequencies and populations. In this way, each MFOP acts as a filter on the ongoing dynamics, allowing connectivity to be resolved by oscillatory state.

For each MFOP, we therefore computed TE as a function of transmission delay *τ* in both directions between populations. This yields a lag-resolved transfer profile that specifies not only whether transfer is boosted, but also when within the relevant oscillatory timescale the enhancement occurs. We then identified TE peaks exceeding surrogate expectations and summarized the corresponding intervals of boosted transfer as simplified diagrams (Fig. 4c,d). We refer to these diagrams as Information Routing Patterns (IRPs), since they represent directed routing between populations. Relative to the effective-connectivity motifs described previously [15, 36], IRPs add an explicit temporal organization: arrows are defined not only by where transfer originates and terminates, but also by when transfer is preferentially expressed. In this sense, they are closer to calling patterns and, more generally, to temporal motifs on temporal networks, where interactions are jointly specified by topology and temporal order [37, 38].

This construction generalizes previous routing analyses in an essential way. In the single-frequency setting of Palmigiano et al. [15], boosted transfer could be described as being gated at specific moments of one oscillatory cycle. Here, by contrast, a given MFOP may involve several coexisting burst components with different frequencies, and the resulting transfer profile may contain multiple temporally separated windows, sometimes in the same direction and sometimes alternating between directions. The relevant routing object is therefore no longer a purely spatial motif of effective connectivity, but a spacetime motif of directed transfer.

Examples from landmark 4 illustrate this clearly. Two MFOPs within the same landmark can differ only in their coordinated burst mask, yet produce markedly different lagged TE profiles and therefore different IRPs (Fig. 4c,d). In one case, boosted transfer is dominated by a single window from the fast to the slow population. In another, additional windows appear, including transfer in the opposite direction and at distinct delays. Thus, even within a fixed landmark configuration, changing the coordinated bursting state is sufficient to reconfigure the temporal pattern of information flow.

Across the full set of MFOPs, this procedure reveals a broad zoo of IRPs (Fig. 4e, derived from full lagged TE profiles shown in Fig. S8). Some are simple, containing a single dominant transfer window in one direction; others comprise several temporally separated transfer events, either repeated in the same direction or distributed across both directions. Yet this diversity is not arbitrary. MFOPs within the same landmark often occupy related temporal windows, suggesting that the structural-dynamical regime defines a coarse temporal scaffold, whereas the specific MFOP selects which transfer events are expressed within that scaffold. Coordinated bursting does not generate routing ex nihilo, but filters and refines a set of structurally permitted communication opportunities.

Several complementary analyses support this interpretation. MFOP-specific coherence, measured relative to periods in which no MFOP is active, is selectively enhanced at the frequencies present in the corresponding burst mask (Fig. S9), indicating that MFOPs reshape the coordination landscape in a frequency-specific manner rather than merely indexing a nonspecific increase in activity. Moreover, MFOPs associated with related IRPs tend to share similar patterns of cross-frequency interaction, including amplitude-amplitude coupling, phase-phase coupling, and lagged cross-correlation structure, although these can differ in strength and asymmetry (Fig. S10, Fig. S11, Fig. S12). These observations suggest that IRPs emerge from a joint interplay of amplitude coordination, phase relationships and inter-population precedence across frequencies, rather than from a single oscillatory mechanism alone.

The multi-frequency character of this routing is essential. When two identical populations are coupled and only one oscillatory timescale is available, the diversity of MFOPs and IRPs is markedly reduced (Fig. S13). Thus, the richness of routing motifs observed here is not a generic consequence of state filtering, but depends specifically on the coexistence and coordination of multiple burst frequencies. Finally, the main routing windows and directions remain qualitatively stable when TE is estimated using richer embeddings of the target past [39], indicating that the inferred IRPs do not depend critically on the simplest embedding choice (Fig. S14).

Together, these results show that MFOPs define oscillatory states with distinct spacetime motifs of directed information transfer. By conditioning connectivity on coordinated multi-frequency bursting states, state filtering reveals how coupling and burst coordination shape when, and in which direction, inter-population communication is selectively boosted. These IRPs provide a mesoscopic description of routing. If that routing is mechanistically meaningful, it should also leave a microscopic trace: MFOPs that favor transfer in a given direction and at specific delays should modulate the probability of spike transmission across the corresponding synaptic pathways. We therefore next ask whether the routing motifs identified by TE are recapitulated at the level of concrete monosynaptic transmission.

### Coordinated multi-frequency bursting selectively reweights delayed spike transmission

The Information Routing Patterns described above identify oscillatory-state-dependent motifs of directed functional connectivity at the mesoscopic level. Yet TE remains a population-level measure, and does not by itself specify how routing is implemented at the level of individual spikes. If the routing motifs associated with coordinated multi-frequency bursting are mechanistically meaningful, they should be accompanied by corresponding changes in the probability that spikes emitted in one population are effectively transmitted to neurons in the other, with a dependence on both transmission direction and effective latency.

This question is non-trivial in a recurrent network. Even when two neurons are linked monosynaptically across populations, the influence of a presynaptic spike need not be confined to its immediate postsynaptic effect. Through recurrent interactions, spikes emitted in the past may reverberate locally before contributing to downstream activation, thereby introducing an effective transmission latency. To capture this possibility without explicitly embedding feedforward synfire-chain motifs into the random microcircuit, we introduced explicit transmission lines with adjustable delay *d* on a subset of excitatory projections between the two populations (Fig. 5a; Methods). This construction provides a parsimonious surrogate for delayed propagation through recurrently embedded transmission motifs [40, 41].

**Figure 5.**
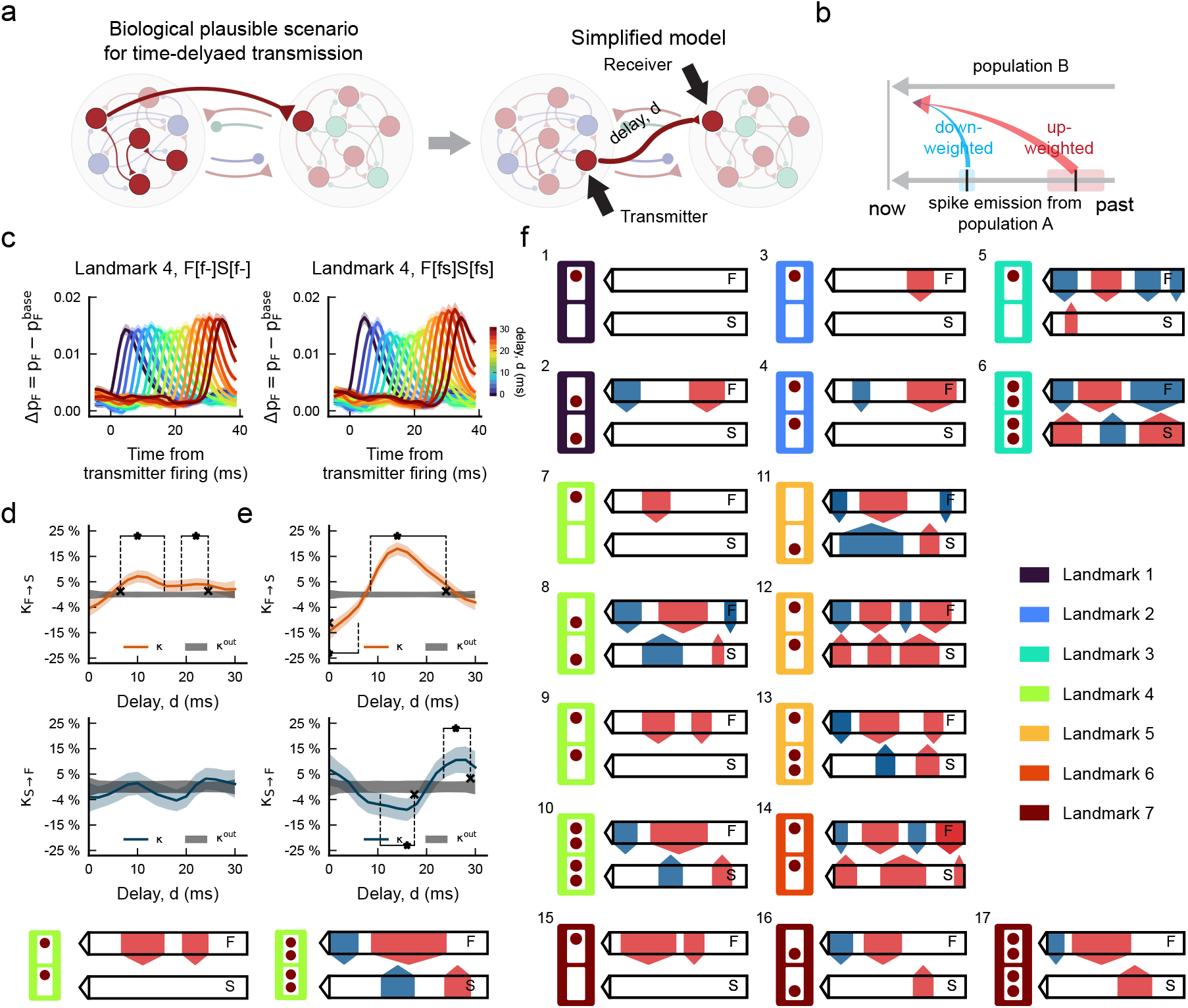
Coordinated multi-frequency bursting selectively reweights delayed spike transmission across monosynaptic pathways. **a**. Conceptual framework for time-delayed spike transmission. Left, biologically plausible scenario in which spikes emitted in one population can influence another after variable effective delays through recurrent interactions. Right, simplified model used here, in which a transmitter neuron projects to a receiver neuron through an explicit transmission line with delay *d*. **b**. Schematic of delay-dependent transmission weighting. For a given direction of transmission, spikes emitted in the past can be either down-weighted or up-weighted depending on their delay relative to the current oscillatory state of the network. **c**. Example receiver responses for two coordinated bursting states from landmark 4. The quantity 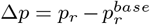 measures the firing probability of the receiver neuron (*p*_*r*_) relative to the baseline probability (*p*_*base*_)of the receiving population *r* ∈ {Pop_*F*_, `Pop`_S_} with the receiver excluded. Curves show responses across transmission delays *d*. In one state, responses vary little with delay; in another state from the same landmark, delayed inputs are selectively suppressed or enhanced over distinct latency ranges. **d**,**e**. Delay-dependent spike-transmission modulation for the two states shown in **c**. Top, modulation *κ* for `Pop`_*F*_ -to-`Pop`_*S*_ transmission; bottom, modulation for `Pop`_*S*_-to-`Pop`_*F*_ transmission. Gray bands indicate the corresponding out-of-state baseline. The shaded region indicates the 5th–95th percentile range. Asterisks mark peaks of significant facilitation or suppression, and crosses delimit the corresponding delay ranges. Bottom schematics summarize the resulting spike-transmission barcode, with red and blue segments denoting up-weighted and down-weighted transmission windows, respectively. **f**. Spike-transmission barcodes identified across coordinated bursting states. Colored frames indicate landmark identity. Each barcode summarizes the direction, delay range and sign of state-dependent modulation in spike transmission across the two populations.

For each coordinated bursting state, we then measured how the firing probability of a receiver neuron varied as a function of this delay. For a given transmission direction, the spike probability of the receiver following delayed input was compared with a baseline firing probability measured in the receiving population while excluding the receiver neuron itself (Fig. 5b,c; Methods). This yielded a delay-dependent modulation profile indicating whether spikes arriving after a given latency were effectively up-weighted or down-weighted under the active state. We summarize this relative modulation by the quantity *κ*, defined as the increase or decrease in spike-transmission probability relative to periods outside the corresponding bursting state.

Examples from landmark 4 illustrate that these modulation profiles are strongly state-dependent. In one state, transmission in the fast-to-slow direction is selectively enhanced over a restricted range of delays, whereas modulation in the opposite direction remains weak (Fig. 5d). In another state from the same landmark, the pattern changes qualitatively: transmission is suppressed at some delays and enhanced at others, and the balance between directions is reorganized (Fig. 5e). Thus, within a fixed structural-dynamical regime, changing the coordinated burst configuration is sufficient to alter the temporal profile of effective spike transfer.

These delay-dependent modulation profiles can be summarized diagrammatically in the same spirit as IRPs, now using red and blue segments to denote up-weighted and down-weighted transmission windows along the delay axis (Fig. 5d–f). Across the full set of states, this procedure reveals a diverse repertoire of spike-transmission barcodes (Fig. 5f; full curves in Fig. S15). Some states preferentially enhance transmission in a single direction and over a narrow delay range, whereas others combine several facilitation and suppression windows, sometimes in both directions. As in the TE analysis, this diversity is structured rather than arbitrary: states within the same landmark often recruit related delay windows, while differing in which of those windows are amplified or attenuated.

This structure is not explained by the mere presence of a fast or slow oscillatory component. If transmission were determined only by which rhythm was active, similar oscillatory content should produce similar delay-dependent modulation. Instead, the same burst components can be associated with different transmission barcodes across landmarks, and different coordinated burst states within a given landmark can likewise generate distinct barcodes despite sharing the same broad oscillatory repertoire. Conversely, when only a single oscillatory timescale is available, as in the control systems with two identical populations, the repertoire of transmission barcodes collapses markedly (Fig. S16). Selective transmission therefore depends on the coordinated multi-frequency state of the circuit, not on the isolated presence of one oscillatory component.

Phase- and frequency-dependent modulation of background spiking probability likely contributes to this effect, but does not fully explain it. Coordinated bursting states reshape the excitability landscape of both populations (Fig. S17), yet the observed barcodes are additionally direction-specific and delay-specific. They therefore reflect not only when neurons are more or less excitable, but which delayed inputs are selectively incorporated into inter-population communication.

Despite being derived from a different observable and at a different scale, these spike-transmission barcodes show a statistically significant similarity to the TE-based IRPs. To quantify this relationship, we compared each state-specific barcode with its corresponding IRP using the Wasserstein distance [42] between the two temporal patterns (Fig. S18). For most states, the observed distance was significantly smaller than expected from surrogate barcodes with randomized arrow positions, indicating that the similarity between mesoscale routing motifs and microscopic transmission patterns is unlikely to arise by chance. The correspondence is not exact, nor should it be expected to be: TE captures directed dependence in collective activity, whereas *κ* measures the modulation of actual delayed spike transmission across monosynaptic pathways. Nevertheless, the two descriptions converge more than expected under a null model, supporting the view that the mesoscale routing motifs identified by TE have a genuine microscopic counterpart.

One systematic difference remains apparent. Spike-based analyses often show an additional enhancement at very short delays, especially under strong synchrony. This likely reflects a generic increase in immediate transmission reliability caused by spike-time alignment, rather than a state-specific routing effect. Because the TE analysis contrasts each oscillatory state against surrogates defined at the population level, such ubiquitous short-latency influences are largely factored out there, whereas they remain visible in the microscopic transmission analysis.

Taken together, these results provide mechanistic support for the routing motifs identified by state-resolved TE. Coordinated multi-frequency bursting does not merely reshape directed dependence between collective signals; it also selectively reweights spike transmission efficacy according to both the direction of propagation and the time at which spikes were emitted in the past. Communication is therefore structured simultaneously at mesoscopic and microscopic scales through transient state-dependent filtering of which delayed inputs are favored and which are suppressed. In this sense, coordinated burst states act as a form of temporal weighting over incoming spike histories, with a functional logic reminiscent of positional weighting or positional encoding in sequence-processing architectures [34]. We will return to the broader implications of this analogy in the Discussion.

## Discussion

Flexible cognition requires communication that is selective not only in space, but also in time. Here we show that such time-structured communication can emerge intrinsically from the interaction of coupled populations generating spontaneous multi-frequency bursts. Three main conclusions follow. First, coupling does not merely juxtapose populations with distinct native oscillatory preferences, but reorganizes them into a structured repertoire of coordinated multi-frequency patterns (MFOPs). Second, these states define Information Routing Patterns (IRPs): spacetime motifs of directed transfer specifying not only who communicates with whom, but also when transfer is preferentially expressed. Third, these mesoscale routing motifs have a microscopic counterpart, in that the same coordinated burst states selectively up-or down-weight delayed spike transmission across monosynaptic pathways. Taken together, these results identify coordinated multi-frequency bursting as a mechanism by which recurrent circuits can transiently filter past inputs according to both source and latency.

Precise inhibitory timing, dendritic nonlinearities, short-term synaptic adaptation, interactions across multiple pathways, and shifts in brain state have all been proposed as substrates for the dynamic control of communication [4–8]. Many of these mechanisms rely on specific structural arrangements or finely tuned local operations. By contrast, the mechanism described here exploits emergent dynamics generated by comparatively weak architectural constraints: modest diversity in inhibitory timescales, recurrent coupling, and stochastic bursting suffice to create a broad repertoire of communication motifs. In this sense, the model emphasizes dynamic degeneracy rather than structural hardwiring. A relatively simple circuit can therefore generate a large family of temporally structured routing states.

Among alternative proposals, traveling waves are particularly relevant because they also have been linked to latency-based reweighting of communication [9]. In wave-based accounts, temporal structure is inherited from spatial propagation: different delays arise because activity moves across an extended substrate. Here, by contrast, both spatial and temporal structure emerge within the dynamics of a compact coupled circuit, without requiring a geometrically extended propagation medium. This does not make the two views mutually exclusive. Rather, it suggests that delay-based routing may be implemented at multiple scales: through long-range spatial propagation in some settings, and through internally generated multi-frequency coordination in others.

Our results also bear directly on influential ideas that assign distinct frequencies to distinct network modules [43, 44], and distinct communication roles to distinct frequency bands [16–18]. In our model, we do not impose a strict anatomical interpretation on the coupled modules, and several readings remain possible. One is inter-areal, in which two nearby cortical regions with distinct resonance frequencies interact through long-range excitation (corresponding to *α* ≫ *β*; cf. Fig. 2a). Another is interlaminar, in which the two populations represent local recurrent circuits in different cortical layers, coupled through reciprocal excitation and inhibition (in which case *β* ≥ *α*). In either interpretation, however, the central point is the same: oscillatory identity is not fixed once and for all by the intrinsic properties of a subpopulation. Coupling can preserve, hybridize, or reassign intrinsic burst preferences. This speaks directly to current debates on interneuronal diversity. PV and SOM circuitry clearly matters [28], but their influence is not reducible to a rigid one-cell-type/one-frequency mapping; rather, both classes contribute in interacting ways to oscillation frequency, amplitude, and stability [29, 45]. Interneuronal diversity does not hardwire but rather enriches the accessible dynamical repertoire and its control possibilities, as further illustrated by the marked collapse of the IRP repertoire when only one interneuronal type is included (Fig. S13).

Whether one adopts an inter-regional or interlaminar perspective, faster rhythms have often been associated with feedforward signalling and slower rhythms with feedback signalling [18–20, 46, 47]. In predictive-coding formulations, this asymmetry is often sharpened further, with faster rhythms linked to bottom-up prediction errors and slower rhythms to top-down predictions [20]. Our results challenge any rigid version of this mapping. In the model, coupling can preserve native frequency contrasts, but more generally it can reassign or even invert them. Both fast and slow components can appear in putatively “upstream” or “downstream” nodes, support communication in either direction, and participate in motifs where the order of influence reverses (for example, IRP #9 versus IRP #4 in Fig. 4). This does not invalidate the empirical observation that certain frequencies may, on average, be enriched in certain directions. It does imply, however, that such averages may conceal a much richer repertoire of transient multi-frequency motifs than is accessible through time-averaged analyses, as already shown in hippocampal or cortical networks [27, 48].

This becomes especially clear when considering MFOPs and IRPs themselves. These states are not simple superpositions of a slow and a fast channel, each endowed with a fixed routing role. The topology of an IRP associated with a combined bursting pattern cannot, in general, be predicted by merely merging the routing effects of simpler motifs: for instance, IRP #10 differs from the plain union of IRPs #8 and #9. Frequency channels therefore do not act independently, but interact synergistically [49, 50], through cross-frequency relationships [51] that vary across landmarks and can be recombined within the same structural regime. Some motifs, such as IRPs #12 and #14, resemble the multi-slot schemes envisioned in theta–gamma models [10] or rhythmic attentional-sampling hypotheses [52]; others, such as IRPs #10 and #6, are closer to frameworks in which slower rhythms gate faster-frequency-mediated communication within specific phase windows [21, 22].

What the present model adds is that these alternatives need not correspond to distinct, fixed circuit regimes. Instead, the same coupled system can transiently instantiate different communication logics depending on which multi-frequency state is active. Thus, a given frequency component does not permanently “mean” feedforward, feedback, or any other routing function. Even identical frequency components observed within the same region can mediate distinct IRPs depending on their coordination with other frequencies (e.g., IRP #9 versus IRP #10 in Fig. 4), while the same set of multi-frequency components can support different routing functions across dynamical regimes (e.g., IRP #9 versus IRP #12). This context dependence is lost when oscillations are considered in isolation (Fig. S13), where similar oscillatory motifs tend to yield similar communication patterns. Together, these findings argue against treating spectral components as independent channels with stereotyped functional roles, and instead call for neural signal analyses that explicitly account for their combinatorial and dynamical coordination.

Once communication is characterized not only by source and target but also by the time at which it is preferentially expressed, the relevant object is no longer a static connectivity motif but a temporal graph [37, 38]. In human phone-call networks, the over-representation of specific temporal motifs reveals latent organization, including homophily, division of roles, and the functional structure of communication chains [31]. A similar interpretive possibility may exist here: IRPs may provide clues about how distributed neural computation is organized in time, or at least about the constraints under which such organization can unfold. They point to communication as a sequentially structured temporal process rather than a collection of isolated events. This suggests a second, equally cautious analogy. As operons coordinate the co-expression of functionally related genes under common control [53, 54], MFOPs may analogously trigger whole families of temporally linked—and perhaps algorithmically chained—communication events. This possibility remains speculative, but it underscores the need for experimental approaches capable of resolving “routing operons,” whenever they exist, rather than isolated communication events or communication on average.

We emphasize that, in our simulations, communication is always evaluated relative to a baseline state in which coordinated transient bursts are absent. What is therefore measured is the gain in communication associated with a specific burst configuration. In this sense, bursts signal communication boosting, supporting a functional rather than merely epiphenomenal role [55]. This does not imply, however, that coherence alone is the unique mechanism underlying the gain. Although coherence is transiently enhanced during MFOPs (Fig. S9), it may be only one signature—together with several forms of cross-frequency coupling (Fig. S10, Fig. S11)—of the circuit entering a state especially suitable for communication. On this view, the boost in communication would arise from broader forms of dynamic coordination rather than from coherence alone.

The microscopic spike-transmission analysis sharpens this picture further. Coordinated multi-frequency bursting selectively reweights spike-transmission efficacy according to both propagation direction and the time at which spikes were emitted in the past. This is the strongest sense in which communication becomes time-structured here: the circuit transiently privileges some delayed inputs while suppressing others. In the present simplified framework, monosynaptic transmission delays are adjusted parametrically to scan a continuous range of possible input latencies. In a more natural setting, one may instead imagine receiving neurons integrating “now” spikes emitted at different times in the past but arriving synchronously because they have propagated along embedded pathways with distinct effective delays [40, 41]. In that sense, the circuit would implement a form of time-to-space transform [56, 57], whereby temporal position in the recent past is converted into distinct channels of arrival in the present. Even if these pathways had equivalent structural efficacy, their impact on the receiver would still be differentially weighted by the coordinated burst state. Functionally, such reweighting is reminiscent of positional weighting or positional encoding in sequence-processing architectures [34]. The analogy must be treated with caution: our model does not represent sequences explicitly as tokenized inputs, nor does it implement learned attention layers. What it does suggest is that a recurrent oscillatory circuit can endogenously assign different weights to incoming spikes according to their recent latency of emission. More speculatively, the diversity of MFOPs may allow one and the same circuit to instantiate several coexisting weighting profiles over recent spike histories, comparable in spirit to the multiple concurrent position-sensitive weighting schemes enabled by multi-head attention [34]. It would therefore be interesting to test whether recurrent networks capable of expressing MFOP-like dynamics gain an advantage on tasks requiring temporal sequence processing or prediction, extending recent proposals of oscillatory machine-learning architectures [58, 59].

We conclude by noting several limitations of the present study. First, the motif catalogue is not exhaustive. We explored only a restricted subset of synchrony levels and coupling parameters, while approximately controlling firing rate; relaxing these constraints would almost certainly enlarge the repertoire further. The aim here is therefore not to enumerate all possible motifs, but to establish the qualitative richness that coordinated multi-frequency bursting can already support. Second, the control of motif occurrence remains open. In the earlier single-frequency setting of [15], spatial biasing could favor particular routing states. Here, selecting among richer spacetime motifs will likely require either structured external forcing or learning rules capable of stabilizing particular MFOPs. Third, the model contains only two populations. Whether similar principles scale to larger circuits and whether motif composition becomes hierarchical in larger networks [60] remains to be established.

These limitations point directly to broader questions. Can larger coupled networks build compositional libraries of communication routines rather than isolated routing events? Can learning regulate not only which motifs exist, but which become accessible under different behavioral contexts? More speculatively, the structured boosting and suppression of delayed spike transmission uncovered here may provide a substrate on which spike-timing-dependent plasticity could act to stabilize alternative transmission motifs, consistent with prior work showing that spike-timing-dependent plasticity can select specific propagation delays during oscillatory activity and organize reproducible delayed spike patterns [61, 62]. In that sense, coordinated bursting may help map temporal structure in the environment onto an internally available repertoire of communication routines.

Our results suggest that spontaneous multi-frequency bursts define coordinated dynamical states that transiently sculpt communication across scales, from directed transfer between collective signals to the selective weighting of delayed spikes. If so, oscillatory bursts may be doing more than routing information: they may provide the brain with an endogenous grammar for temporally structured computation.

## Methods

### Single-population model

We considered two single-population network models, a fast population (`Pop`_*F*_) and a slow population (`Pop`_*S*_). Each population contained 800 excitatory (*E*) and 200 inhibitory (*I*) neuron models [63], preserving the 4:1 excitatory-to-inhibitory ratio commonly observed in cortex [64]. The single-neuron dynamics were identical across all neuron classes. The only difference between `Pop`_*F*_ and `Pop`_*S*_ lay in their inhibitory synapses: inhibitory synapses in `Pop`_*S*_ had slower rise and decay time constants than those in `Pop`_*F*_. This simplification was motivated by physiological evidence that distinct interneuron subtypes contribute differently to cortical network regulation and oscillatory dynamics. In particular, parvalbumin-positive interneurons have been associated with gamma-range oscillations, whereas somatostatin-positive interneurons have been linked to beta or slow-gamma oscillations [29, 30]. Although these interneuron classes differ in many physiological respects, here we reduced this diversity to two inhibitory synaptic types.

For a presynaptic neuron of type *x* ∈ {*E*_*F*_, *E*_*S*_, *I*_*F*_, *I*_*S*_}, the synaptic conductance was modeled as proportional to the channel opening probability *P*_*x*_(*t*), which followed a double-exponential form [65]:

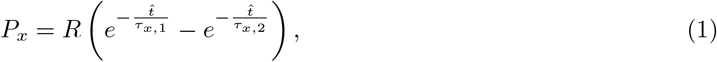

where 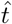 denotes the time elapsed since the last spike of the presynaptic neuron, and *R* is a normalization constant. The parameters *τ*_*x*,1_ and *τ*_*x*,2_ determine the rise and decay time constants, with

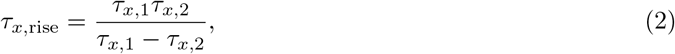

and

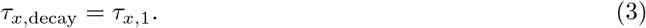

The values used for each neuronal type are summarized in Table 1.

**Table 1:** Synaptic time constants used in the simulations.

| | $E_F$ | $E_S$ | $I_F$ | $I_S$ |
| --- | --- | --- | --- | --- |
| $\tau_{x,1}$ | 1 | 1 | 2.5 | 8 |
| $\tau_{x,2}$ | 0.3 | 0.3 | 0.5 | 1 |

Excitatory neurons in both populations shared the same synaptic time constants, with faster rise and decay than inhibitory neurons [66]. By contrast, inhibitory neurons differed across the two populations, with `Pop`_*S*_ having the longest inhibitory time constants. This difference allowed `Pop`_*S*_ to exhibit intrinsically slower dynamics than `Pop`_*F*_ at comparable levels of network synchrony.

The postsynaptic current was modeled as

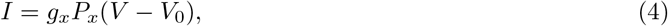

where *g*_*x*_ denotes the maximal synaptic conductance, *V* is the postsynaptic membrane potential, and *V*_0_ is the synaptic reversal potential, with *V*_0_ = 0 mV for excitatory presynaptic neurons and *V*_0_ = − 80 mV for inhibitory presynaptic neurons.

Connections within each population were random, with probabilities depending only on the source neuron type. For example, an *E*_*F*_ neuron projected to both *E*_*F*_ and *I*_*F*_ neurons with probability *p*_*E*_. Each neuron also received background excitatory synaptic input modeled as a Poisson process with rate *ν*. These inputs induced synaptic currents with maximal conductance *g*_*P*_ = 0.002, and their rate *ν* was population-specific. Unless otherwise stated, all simulations assumed zero synaptic delay (*t*_*d*_ = 0 ms).

Within each population, the excitatory projection probability *p*_*E*_ was taken to scale linearly with the inhibitory projection probability,

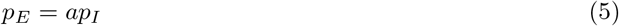

where 0 ≤ *a* ≤ 1 reflects the relatively dense local projection of inhibitory interneurons [67]. The inhibitory conductance scaled as

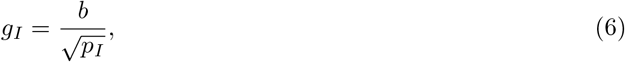

and the excitatory conductance as

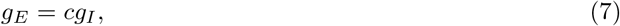

On the iso-firing rate line, the background input rate was followed as

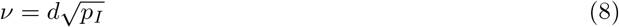

consistent with formulations used to obtain balanced excitation–inhibition states [68, 69]. The specific parameter values are given below.

Simulations were run for 30 s using a fourth-order Runge–Kutta method with a time step of 0.05 ms. We verified that further reducing the time step did not alter the dynamics. To reduce stochastic variability, each parameter condition was simulated across 200 independent trials with different random seeds.

### Characterizing single-population dynamics

To characterize the single-population dynamics and determine the operating regime, we quantified synchrony, firing rate, and spike-time irregularity.

Synchrony within a population was quantified by *χ* [69], defined as

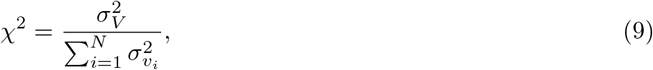

where 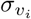 denotes the standard deviation of the membrane potential of neuron *i*, and *σ*_*V*_ is the standard deviation of the population-averaged membrane potential

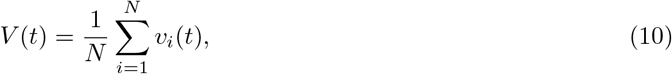

with *N* the number of neurons. Here *V* (*t*) was taken as an LFP-like signal. In the fully synchronized regime, all neurons fluctuate coherently and *χ* → 1. In the asynchronous regime, uncorrelated fluctuations largely cancel in the population average, and *χ* decreases as 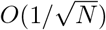 because of finite-size effects [69].

The population firing rate was defined as the average number of spikes per neuron over the simulation, excluding the first 0.5 s. Neural activity *in vivo* rarely displays fully synchronized firing. Instead, it is typically irregular, which we quantified using the coefficient of variation of inter-spike intervals, *CV*_*ISI*_ :

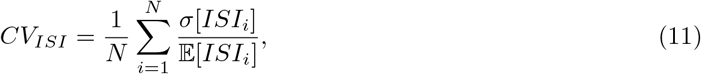

where *σ*[*ISI*_*i*_] and 𝔼 [*ISI*_*i*_] denote the standard deviation and mean of the inter-spike intervals of neuron *i*, respectively. In balanced excitation–inhibition regimes, *CV*_*ISI*_ approaches 1, reflecting Poisson-like firing; similarly, neuronal activity *in vivo* often shows *CV*_*ISI*_ values close to 1 [70]. Values larger than 1 indicate highly irregular firing, whereas 0 corresponds to perfectly regular firing. In our simulations, *CV*_*ISI*_ in both `Pop`_*F*_ and `Pop`_*S*_ varied from approximately 0.6 to 0.8, indicating irregular, although not maximally irregular, activity.

To isolate the effect of synchrony from changes in overall excitability, we identified an iso-firing-rate line by a grid search over the local parameter space of each population. Along this line, both populations maintained similar firing rates and matched synchrony levels, while synchrony itself could vary. Specifically, *χ* ranged from 0.1 to 0.4 in both populations, while the firing rate remained close to 8 Hz. For larger *χ*, firing rates increased substantially (see Fig. 2c,d), so the iso-firing-rate line could not be extended further while preserving the target firing-rate regime. The corresponding parameter values are listed in Table 2, with *p*_*I*_ ranging from 0.153 to 0.702 in `Pop`_*F*_ and from 0.07 to 0.585 in `Pop`_*S*_. We defined *echelon* as the position along this iso-firing-rate line, ranging from 0 to 1, and simulated the activity at echelon values 0, 0.5, and 1. The resulting changes in *χ*, firing rate, and *CV*_*ISI*_ are shown in Fig. S1a.

**Table 2:**
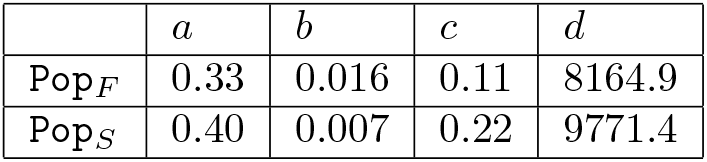
Parameters used to define local projection probabilities, conductances, and background input rates in the fast and slow populations.

| | $a$ | $b$ | $c$ | $d$ |
| --- | --- | --- | --- | --- |
| $\text{Pop}_F$ | 0.33 | 0.016 | 0.11 | 8164.9 |
| $\text{Pop}_S$ | 0.40 | 0.007 | 0.22 | 9771.4 |

### Inter-population connectivity

To study communication through multi-frequency oscillations, we introduced bidirectional coupling between `Pop`_*F*_ and `Pop`_*S*_. These populations can be interpreted as distinct subpopulations within one brain region, as closely connected regions, or as different cortical layers. Inter-population connectivity therefore included both excitatory and inhibitory projections, with possible directional asymmetry.

We parameterized the inter-population connectivity by three structural parameters: the inter-excitatory projection ratio *α* ∈ [0, 2], the inter-inhibitory projection ratio *β* ∈ [0, 1], and an asymmetry factor *ω*∈ [− 1, 1]. The asymmetry factor specifies how excitatory and inhibitory projections are distributed between the two populations. We defined the directional scaling factors

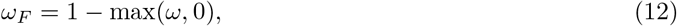

and

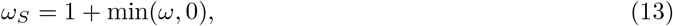

such that *ω >* 0 weakens projections from the `Pop`_*F*_ to `Pop`_*S*_, whereas *ω <* 0 weakens projections from the `Pop`_*S*_ to `Pop`_*F*_. When *ω* = 0, the inter-population connectivity is symmetric.

The inter-population excitatory projection probabilities were defined as

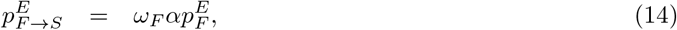

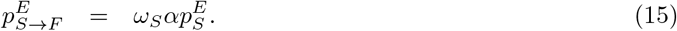

Similarly, the inter-population inhibitory projection probabilities were

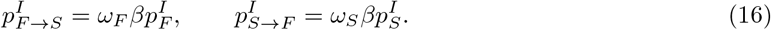

Here, *α >* 1 indicates that excitatory projection between the two populations is stronger than the corresponding within population excitatory projection, whereas *α <* 1 reflects weaker inter-population exctiatory projection. In contrast, *β* was restricted to the range [0, 1] so that inter-population inhibitory projections remained at most as dense as the corresponding within population inhibitory projection.

For the control condition with two identical populations (Fig. S13 and Fig. S16), the same inter-population connectivity was used, but with 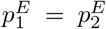 and 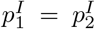, where 1 and 2 denote the two populations. The ranges of *α* and *β* were unchanged, whereas *ω* was restricted to [0, 1] because the two populations were identical.

In all cases, these projections targeted both excitatory and inhibitory neurons in the recipient population. The maximal synaptic conductances of inter-population projections were set equal to those of the corresponding local projections, based on the source population parameters.

### Characterizing multi-frequency oscillatory dynamics

We next characterized the oscillatory dynamics that emerged when `Pop`_*F*_ and `Pop`_*S*_ were interconnected. These dynamics varied as a function of the structural parameters (*echelon, α, β, ω*). To describe these regimes, we defined a set of dynamical features summarizing local oscillatory activity and inter-population coordination. These features were computed from the LFP-like signals *V*_*F*_ and *V*_*S*_ of `Pop`_*F*_ and `Pop`_*S*_, respectively. From these signals we calculated both within-population autocorrelations and between-population cross-correlations. The dynamical features are summarized in Table 3. All features except *C*^*M*^ and *τ*^*C*^ were computed separately for each population.

**Table 3:** Dynamical features used to characterize oscillatory structure and inter-population coordination.

| Dynamical feature | Description |
| --- | --- |
| $AC^1$ | First non-zero local maximum of the autocorrelation, excluding lag 0 |
| $AC^M$ | Global maximum of the autocorrelation, excluding lag 0 |
| $\tau^1$ | Time lag at which $AC^1$ occurs |
| $\tau^M$ | Time lag at which $AC^M$ occurs |
| $C^M$ | Maximum value of the cross-correlation between the LFP-like signals of the two populations |
| $\tau^C$ | Time lag at which $C^M$ occurs in the cross-correlation |

The autocorrelation amplitudes *AC*^1^ and *AC*^*M*^ quantify the strength of, respectively, the fastest oscillatory component and the dominant oscillatory component. Both range from 0 to 1, with larger values indicating stronger rhythmic activity. The corresponding time lags *τ*^1^ and *τ*^*M*^ characterize the associated timescales of the LFP-like fluctuations: *τ*^1^ captures the fastest oscillatory timescale, whereas *τ*^*M*^ captures the timescale of the dominant oscillation. When the autocorrelation only shows a single local maximum (excluding lag 0), we set *AC*^1^ = *AC*^*M*^ and *τ*^1^ = *τ*^*M*^. These values can be interpreted as inverse frequencies by the convolution theorem [71]. Finally, *C*^*M*^ and *τ*^*C*^ quantify temporal coordination between the two populations. *C*^*M*^ measures the strength of inter-population coordination, normalized between 0 and 1, and *τ*^*C*^ indicates the corresponding lead–lag relationship. Positive *τ*^*C*^ indicates that *V*_*F*_ lags behind *V*_*S*_, whereas negative *τ*^*C*^ indicates the opposite. The extraction of these features is illustrated in Fig. S2.

These quantities were computed iteratively in overlapping 1-s windows across the entire simulation, excluding the first 1 s to remove initialization transients. For each simulation, we retained both the mean and the variation of each quantity in order to capture not only the average oscillatory activity but also its temporal variability under transient bursting. The resulting dynamical feature vector 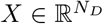 had dimension *N*_*D*_ = 20, corresponding to the mean and variation of the 10 feature quantities.

The feature vector *X* was determined by the structural parameters (*echelon, α, β, ω*). To visualize how these features varied across structural configurations, we applied uniform manifold approximation and projection (UMAP) to the set 𝕏 = *X*_(*echelon,α,β,ω*)_. UMAP provides an interpretable low-dimensional representation of the feature space by approximately preserving its topological structure. The embedding was computed using the Python library [72] with `n_neighbors`=200, `min_dist`=0.8, `n_components`=2, and Euclidean distance as the metric. The full structural and dynamical parameters projected onto the UMAP space are shown in Fig. S3.

### Clustering analysis

Multiple structural parameters produced similar oscillatory dynamics. To group simulations according to their emergent oscillatory dynamics rather than pre-selected parameters, we applied unsupervised clustering to the dynamical feature matrix 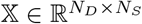, where each column corresponds to the feature vector of one simulated sample, and *N*_*S*_ is the number of simulated parameter sets.

We used consensus clustering [73], an ensemble-based method that aggregates multiple runs of K-means clustering with different initializations. For each value of *K*, we ran K-means *N*_*C*_ times with different random seeds and constructed a consensus matrix 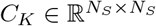, where each element [*C*_*K*_]_*ij*_ is the proportion of runs in which samples *i* and *j* were assigned to the same cluster:

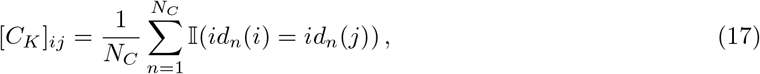

where *id*_*n*_(*i*) denotes the cluster label of sample *i* in the *n*th run, and 𝕀(·) is the indicator function.

This procedure was repeated for *K* = 1, …, 10, and the final number of clusters was selected using two criteria. First, we identified the local maximum of the second derivative of the K-means inertia curve, following the elbow criterion. Second, we computed the proportion of ambiguous clustering (PAC), defined as the fraction of sample pairs for which 0.1 *<* [*C*_*K*_]_*ij*_ *<* 0.9, and selected the local minimum of the PAC curve [73]. Both criteria independently yielded *K*^*^ = 7 (see Fig. S4(a)).

After selecting *K*^*^, we performed agglomerative hierarchical clustering [74] using the distance matrix

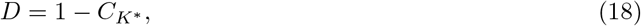

to obtain the final cluster assignments. Fig. S4(b) shows the reordered distance matrix and the corresponding hierarchical linkage. Each resulting cluster was referred to as a distinct dynamical regime. The organization of the dynamical features within each regime is shown in Fig. S4(c), and the distribution of regimes across parameter space in Fig. S4(d).

To identify representative examples within each regime, we computed the silhouette score of every sample after clustering [75], and selected as landmark the sample with the highest score. The corresponding landmark parameters are listed in Table 4.

**Table 4:** Landmark structural parameters for the seven dynamical regimes.

| Regime ID | Echelon | $\alpha$ | $\beta$ | $\omega$ |
| --- | --- | --- | --- | --- |
| 1 | 0 | 1.29 | 0.50 | 0.85 |
| 2 | 0 | 1.00 | 0.78 | -0.30 |
| 3 | 1 | 1.28 | 0.71 | 0.90 |
| 4 | 0.5 | 1.14 | 0.29 | 0.15 |
| 5 | 1 | 0.85 | 0.50 | -0.30 |
| 6 | 1 | 0.85 | 0.85 | -0.30 |
| 7 | 1 | 0.57 | 0.14 | 0.15 |

### Detecting transient oscillations and multi-frequency oscillatory patterns

Transient oscillatory epochs were detected using the procedure described in [27]. For each population, we computed the power spectrogram of the LFP-like signal *V* (*t*) using the short-time Fourier transform (STFT). To detect transient events, we first binarized the spectrogram at a threshold of *m* + 5*σ*, where *m* and *σ* are the mean and standard deviation of the power over time. This isolates sparse blobs corresponding to the most prominent oscillatory bursts. The threshold was then iteratively lowered until it reached *m* + 1.96*σ*, corresponding to a 95% confidence interval under a Gaussian assumption. At each threshold, newly appearing or expanding connected power blobs were tracked across thresholds. At the end of the procedure, isolated blobs were retained as candidate transient oscillatory events. We excluded blobs whose duration was shorter than three cycles of the corresponding oscillation frequency.

While this method preserves both the duration and center frequency of transient oscillations, it can also detect spurious events when fluctuations approach the background level. This occurred more often in interconnected populations, where noise and correlated interactions created null oscillation events. To limit this effect, we constrained the detectable frequency range using the underlying spectral structure. Specifically, the power spectrum of each population was approximated by either one or the sum of two Cauchy distributions. We fitted the spectrum accordingly and extracted the frequency bands corresponding to the half-width at half-maximum of each fitted peak. These bands were then used as frequency bounds for burst detection.

When the two populations were interconnected, transient oscillations in different frequency bands often co-occurred or were temporally coupled (see Fig. S6). Based on the possible combinations of frequency bands and source populations, there were 15 possible bursting states. However, the corresponding oscillatory epochs were not always perfectly overlapping. We therefore identified a common bursting state whenever the overlap duration between two or more transient oscillations exceeded half the duration of each participating oscillation. The corresponding epoch was then defined by the minimum and maximum time points across the overlapping bursts. This allowed us to group closely co-occurring bursts into a multi-frequency oscillatory pattern (MFOP). MFOP occurrence rates are shown in Fig. S7.

Epochs outside any detected transient oscillatory event were labeled as out-of-MFOP periods. These periods do not correspond to an active suppression of oscillations relative to baseline, but rather to times at which activity remained below the burst-detection threshold.

### Cross-frequency coupling and spike-phase coordination analysis

To further characterize the oscillatory interactions associated with each MFOP, we quantified three forms of coupling: phase-phase coupling, amplitude-amplitude coupling, and spike-phase coordination. These measures were computed for all intra- and inter-population frequency pairs (*f*_*X*_, *f*_*Y*_), with *X, Y* ∈ {`Pop`_*F*_, `Pop`_*S*_}, thus capturing both same-frequency and cross-frequency interactions.

For a given population *X* and center frequency *f*_*X*_, we extracted the instantaneous phase and amplitude of the LFP-like signal in the band [*f*_*X*_ − Δ*f, f*_*X*_ +Δ*f* ] using the Hilbert transform. These quantities are denoted by 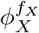 and 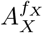, respectively.

To compute phase-phase coupling (PPC) for a frequency pair (*f*_*X*_, *f*_*Y*_), samples of 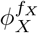 and 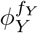 were collected over the time points belonging to a given MFOP and binned into *n*_*f*_ equal bins over [0, 2*π*). Statistical dependence between the two instantaneous phases was quantified using their mutual information:

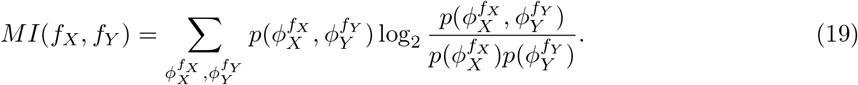

PPC for each MFOP and frequency pair (*f*_*X*_, *f*_*Y*_) was then defined as the normalized mutual information,

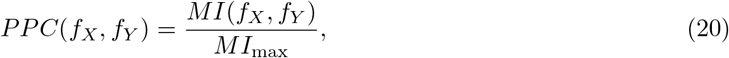

where *MI*_max_ = log_2_(*n*_*f*_) is the maximum possible mutual information for *n*_*f*_ bins. Thus, *PPC*(*f*_*X*_, *f*_*Y*_) approaches 0 when the two phases are statistically independent and approaches 1 when their relationship is maximally constrained.

Amplitude-amplitude coupling (AAC) was quantified analogously, using the binned amplitudes 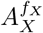 and 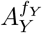 in place of the instantaneous phases. For both PPC and AAC analyses, we used Δ*f* = 2 Hz and *n*_*f*_ = 31. The resulting PPC and AAC values for each MFOP are shown in Fig. S11 and Fig. S10, respectively.

Spike-phase coordination was quantified as the firing probability of one population as a function of the instantaneous phase of the other population. For each MFOP and each frequency band, spike times from one population were assigned to the phase bins of the other population, yielding a phase-dependent firing probability distribution. The resulting spike-phase coordination profiles are shown in Fig. S17.

### Transfer entropy analysis

Transfer entropy (TE) is a nonparametric information-theoretic measure of directed information exchange between two dynamical systems *X* and *Y* [35]. Because TE makes no assumption of linearity or specific temporal structure, it is well suited to the analysis of complex oscillatory signals.

Let *X* = {*x*_*t*_} _*t*=1,…,*T*_ and *Y* = {*y*_*t*_} _*t*=1,…,*T*_ be stationary time series measured simultaneously from two systems. TE quantifies the deviation from the generalized Markov property

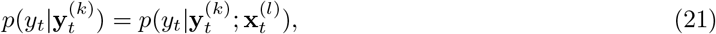

where *p*(·) denotes a transition probability, and 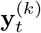 and 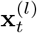 are the past histories of length *k* and *l*, respectively. If the deviation is negligible, the past of *X* provides no additional information about the transition probabilities of *Y*. Conversely, a substantial deviation indicates that the Markov property of *Y* is broken by influences from *X*.

This deviation is quantified by the Kullback–Leibler divergence between the conditional probabilities 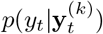 and 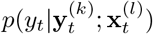:

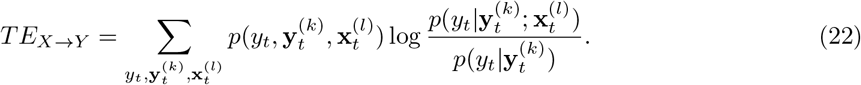

For the history processes 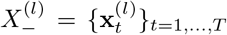 and 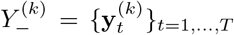, this is equivalent to the conditional mutual information

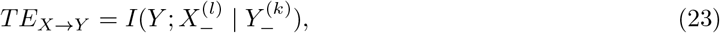

that is, the information shared between the current state of *Y* and the past of *X*, conditioned on the past of *Y* .

To explicitly incorporate interaction delays, we relaxed the history vectors and replaced 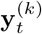 and 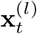 by single time points *y*_*t*−*s*_ and *x*_*t*−*τ*_, as commonly done in delay-resolved TE analyses [15, 36]. For the lagged signal sets *X*_−*τ*_ = {*x*_*t*−*τ*_ }_*t*=*τ*+1,…,*T*_ and *Y*_−*s*_ = {*y*_*t*−*s*_}_*t*=*s*+1,…,*T*_, TE becomes

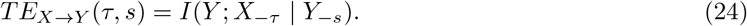

A common choice is to set *τ* = *s*, but this can lead to spurious estimates of directed information transfer when the two systems differ in their autocorrelation structure [76]. To reduce sensitivity to this effect, we computed a delay-specific transfer entropy by averaging over conditioning lags *s*:

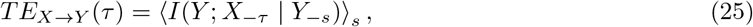

where ⟨·⟩_*s*_ denotes averaging over *s*. Intuitively, this procedure reduces the contribution of effects tied to the autocorrelation of *Y* at any particular lag. In practice, *s* was averaged over one period of the slow oscillation, under the assumption that the structure of information transmission is approximately redundant across slow cycles.

Because conditioning on a single lag may nonetheless remain insufficient for oscillatory signals with richer autocorrelation structure, we additionally estimated transfer entropy using a multi-point embedding of the target past [39, 77]. Specifically, the single conditioning term *Y*_−*s*_ was replaced by an *n*-dimensional history vector,

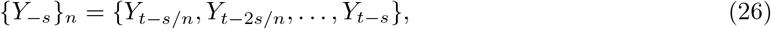

which spans the same temporal interval as the single-lag conditioning term, with ∥{*Y*_−*s*_ }∥_*n*_ = *n* (see also [15]). As shown in Fig. S14, the resulting IRPs were quantitatively similar to those obtained with single-lag conditioning, indicating that the main TE peaks were not driven by bias associated with any specific conditioning lag.

TE was estimated from MUA-like population activity (spike counts per time bin) extracted from the fast and slow populations. Estimation relied on a Gaussian-copula-based entropy estimator implemented in the Python package `Frites` [78], which is well suited to continuous neural signals. To reveal state-dependent changes in information transmission, TE was computed from the time series 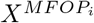 and 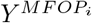 restricted to samples belonging to MFOP *i*.

Oscillatory signals with strong amplitude modulation or complex phase structure can produce non-zero TE even in the absence of true information transmission. To control for this, we constructed surrogate datasets using the iterative amplitude-adjusted Fourier transform (IAAFT) procedure [79]. This method preserves each signal’s amplitude distribution and phase structure while disrupting cross-signal dependencies. TE computed from these surrogates (*TE*_*surr*_) therefore reflects the TE expected from oscillatory structure and autocorrelation alone. True TE was considered as significant only when its median exceeded the 95-th percentile of the surrogate distribution.

However, significance alone was not sufficient to define an IRP component. Transfer entropy could remain above the surrogate level over broad delay ranges, including local troughs between neighboring peaks. Such delays indicate non-zero information transfer, but not necessarily a locally enhanced transfer regime. Because our aim was to summarize delay ranges in which transfer was preferentially boosted, we defined IRP components in a peak-centered manner.

After identifying a significant TE peak at delay *τ*_peak_ (asterisk symbol in Fig. 4c,d), we estimated its temporal extent as follows. Starting from the peak, we first identified the nearest delay on one side (cross symbol in Fig. 4c,d) at which TE fell to the 95th percentile of *TE*_surr_. We then used the TE value at this threshold crossing to identify the matching delay on the opposite side of the peak. These two delays defined the range of the IRP component, which therefore always contained the significant TE peak.

An IRP component should therefore be interpreted as a conservative, peak-centered extent of a TE feature, rather than as the exact interval over which TE exceeds the surrogate threshold. The full IRP was then defined as the collection of all such components across MFOPs, thereby encoding both the delay range and the direction of each peak. The diagrams in the bottom row of Fig. 4c,d provide a compact summary of the IRPs, where width denotes temporal extent and arrowhead direction denotes the direction of information transfer.

By comparing IRPs across MFOPs, we identified how multi-frequency oscillatory dynamics reshape the temporal structure of information flow.

### Quantifying the effect of MFOPs on delayed spike transmission

The effect of MFOPs on delayed spike transmission was examined by selectively modifying the synaptic delays of a small subset of excitatory inter-population projections. Specifically, we selected small sets of transmitter and receiver neurons to probe the effect of conduction delay on spike transfer. To isolate this effect, transmitter–receiver pairs were assigned randomly such that each receiver neuron received inter-population excitatory input exclusively from its paired transmitter neuron. We used ten transmitter and ten receiver neurons in each population, a number small enough not to alter the overall network dynamics. This procedure followed the strategy of [36].

Receiver neurons had the same connectivity as other neurons in their own population, except for the inter-population excitatory input. Each receiver received excitatory input solely from its paired transmitter. Transmitter neurons also retained their usual connectivity, except that each transmitter formed a tenfold-strengthened synaptic projection onto its paired receiver neuron, with a conduction delay *d* sampled between 0 and 30 ms in 2-ms intervals. This strengthened projection compensated for the removed inter-population excitatory inputs to the receiver and emphasized the effect of conduction delay on spike transmission. Because transmitter neurons still fired according to the intrinsic population dynamics, this setup allowed us to track how MFOPs modulated delay-dependent spike transmission under otherwise natural network activity.

For population *r* ∈ {`Pop`_*F*_, `Pop`_*S*_ }, the firing-rate response of each receiver neuron, *p*_*r*_(Δ*t, d*), was aligned to the spike times of its paired transmitter as a function of conduction delay *d* and time relative to the transmitter spike, Δ*t*. However, oscillatory population activity can modulate the firing of receiver neurons independently of the imposed transmission delay. To isolate the specific contribution of delayed transmission, we computed a baseline profile 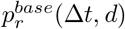 by aligning the responses of non-receiver neurons to the same transmitter spikes at the same delay *d*. This baseline captures oscillation-driven modulation unrelated to the modified conduction delay. The delay-specific response was therefore defined as

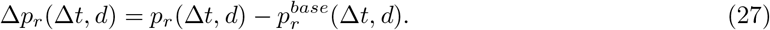

The effect of MFOPs on delayed spike transmission was then quantified relative to out-of-MFOP periods. For transmission from population *X* to population *Y* (*X, Y* ∈ {`Pop`_*F*_, `Pop`_*S*_}), we defined

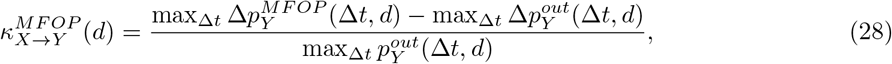

where 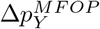 was computed using transmitter spikes occurring during the intervals corresponding to a given MFOP, and 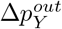 using transmitter spikes occurring outside MFOPs. The quantity 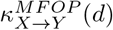 ranges from − 1 to 1, and quantifies the extent to which spike transmission at delay *d* is facilitated (up-weighted) or suppressed (down-weighted) during a given MFOP relative to out-of-MFOP periods. It therefore summarizes how multi-frequency oscillatory dynamics reshape the temporal windows of spike transmission as a function of conduction delay.

The spike transmission pattern was then quantified analogously to the IRP, but using *κ* and taking the out-of-MFOP profile, *κ*^*out*^, rather than surrogate TE, as the reference. Specifically, we first identified the local extrema (maxima and minima) of 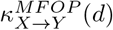. For each extremum (asterisk symbol in Fig. 5d,e), we defined a component as the range of delays over which 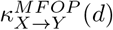 had the same sign as the extremum and satisfied 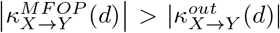, where the reference value on the right-hand side was taken at the nearest extremum of opposite sign (cross symbol in Fig. 5d,e). The resulting components were summarized in a diagram analogous to the IRP (bottom row of Fig. 5d,e), in which width indicates the delay range and arrowhead direction indicates the direction of transmission. Because both local maxima and minima were included, components were additionally color-coded: red for up-weighting (positive *κ*, maxima) and blue for down-weighting (negative *κ*, minima).

## Supporting information

Supplementary Informations

## Code availability

All code required to perform the simulations, extract MFOPs and their associated IRPs, and analyze the modulation of spiking transmission probability is available at: https://github.com/jyKim-97/cross-frequency-burst-routing.

