## Supplementary Informations for "Time-structured communication through cross-frequency bursts"

**This PDF file includes:**

1. Figure S1
2. Figure S2
3. Figure S3
4. Figure S4
5. Figure S5
6. Figure S6
7. Figure S7
8. Figure S8
9. Figure S9
10. Figure S10
11. Figure S11
12. Figure S12
13. Figure S13
14. Figure S14
15. Figure S15
16. Figure S16
17. Figure S17
18. Figure S18

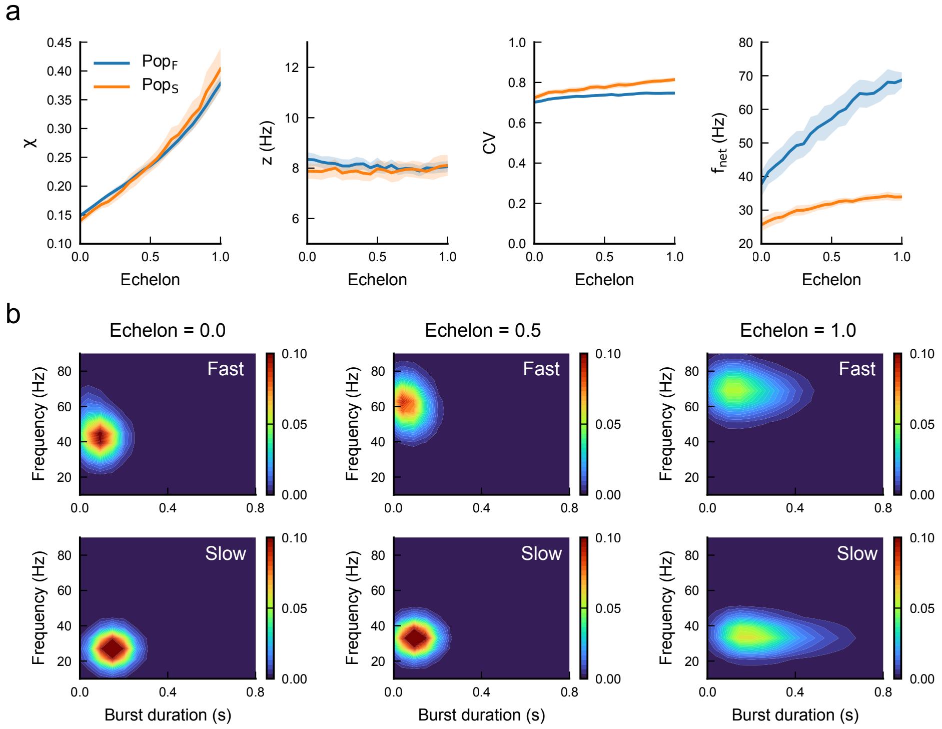


**Figure S1: Iso-firing-rate trajectories vary synchrony while preserving burst-like activity. a**, Synchrony level ($\chi$), firing rate ($z$), coefficient of variation of inter-spike intervals ($CV_{ISI}$), and network frequency ($f_{net}$) measured along the iso-firing-rate trajectory for $\texttt{Pop}_F$ (blue) and $\texttt{Pop}_S$ (orange) populations. Increasing echelon raises synchrony while maintaining approximately constant firing rate. Network frequency remains consistently higher in $\texttt{Pop}_F$ than in $\texttt{Pop}_S$. Shaded regions indicate variability across simulations. $CV_{ISI}$ was computed from the inter-spike intervals of individual neurons (Methods). **b**, Burst-feature densities for the fast (top) and slow (bottom) populations at echelon = 0, 0.5 and 1. Increasing echelon strengthens burst structure and shifts burst frequencies upward, while activity remains transient in both populations across the full range of conditions.

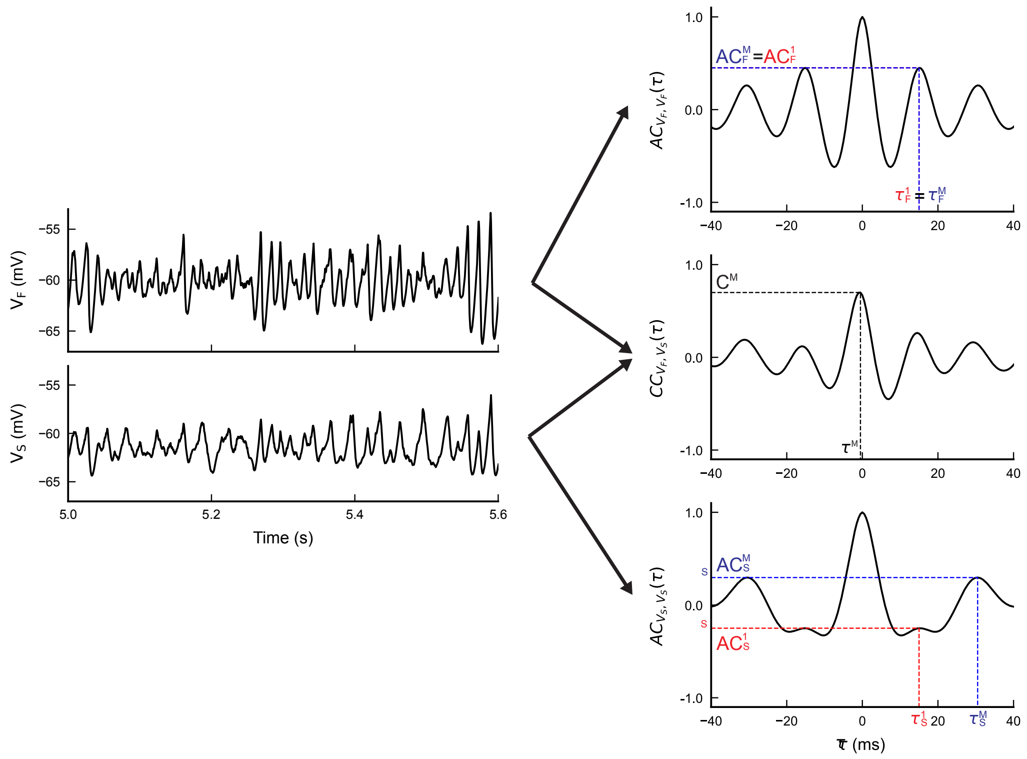


**Figure S2: Extraction of oscillation-related dynamical features from coupled population activity.** Left, example LFP-like signals from the $\texttt{Pop}_F$ ($V_F$) and $\texttt{Pop}_S$ ($V_S$). Right, corresponding auto-correlations for each population and cross-correlation between populations as functions of time lag $\tau$. For each auto-correlation, $AC^M$ denotes the largest non-zero-lag peak and $\tau^M$ its corresponding lag, whereas $AC^1$ and $\tau^1$ denote the first non-zero-lag peak closest to $\tau = 0$. In here, $\texttt{Pop}_F$ shows single frequency oscillation ($AC_F^M = AC_F^1$ and $\tau_F^1 = \tau_F^M$) while $\texttt{Pop}_S$ exhibits both slow and fast frequency oscillations. For the cross-correlation, $C^M$ denotes the maximum value and $\tau^C$ the lag at which it occurs. These quantities contribute to the oscillation-related feature set used to characterize coupled dynamical regimes (Methods).

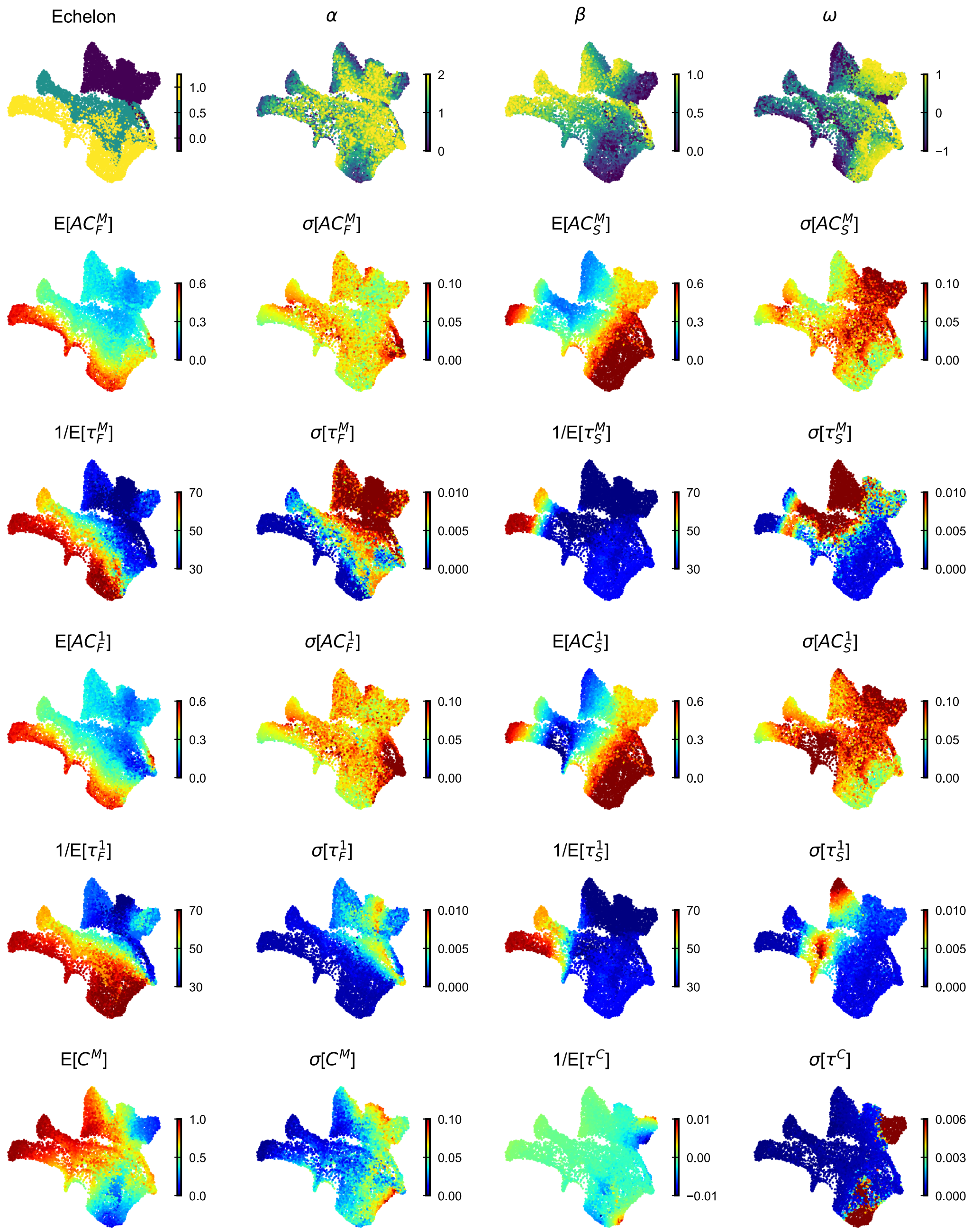


**Figure S3: Structural parameters and oscillation-related dynamical features across the UMAP embedding.** The UMAP embedding was computed from the full oscillation-related feature set. Each point corresponds to one coupled-circuit configuration. Top row, structural parameters projected onto the embedding: echelon, $\alpha$, $\beta$ and $\omega$. Remaining panels show the mean and trial-to-trial variability of the dynamical features extracted from the $\texttt{Pop}_F$ and $\texttt{Pop}_S$, as well as from their cross-correlation structure. Subscripts $F$ and $S$ denote the population from which each feature was measured, and $\mathbb{E}$ and $\sigma$ indicate mean and standard deviation across trials. See Methods for detailed definitions of the parameters.

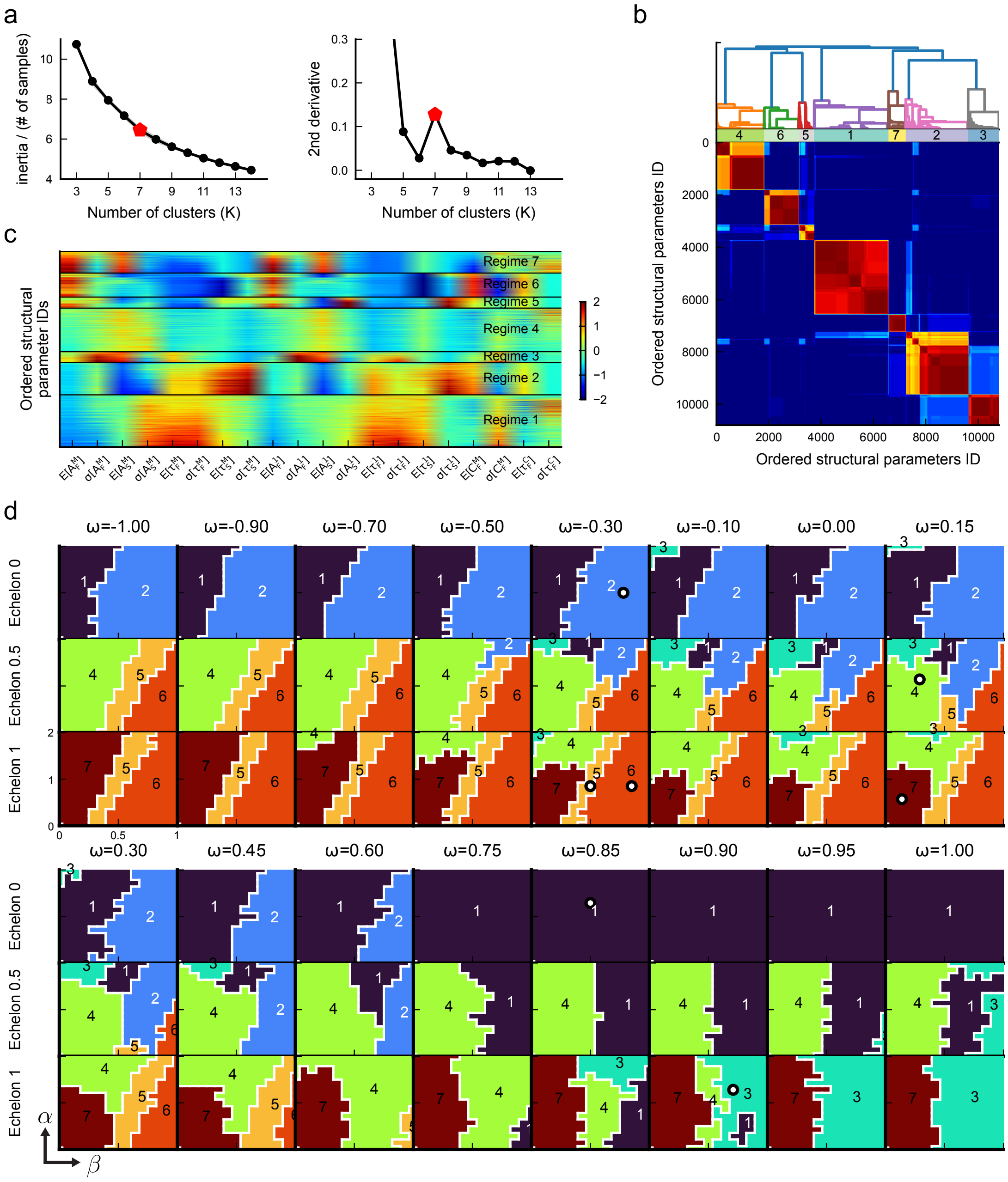

**Figure S4: Unsupervised clustering identifies seven recurrent dynamical regimes in the coupled system.** **a**, K-means inertia (left) and its second derivative (right) as functions of the number of clusters $K$. The peak in the second derivative identifies $K = 7$ as the selected partition. **b**, Consensus clustering matrix after hierarchical reordering. Matrix entry $c_{ij}$ measures how often structural parameter sets $i$ and $j$ were assigned to the same cluster across repeated runs; colors range from low to high consensus. The dendrogram indicates hierarchical relations among the reordered samples. **c**, Oscillation-related dynamical features for all samples grouped by regime and ordered by silhouette value within each regime. Each regime is associated with a characteristic pattern of burst dynamics. **d**, Spatial distribution of dynamical regimes across structural parameter space and corresponding landmarks (○). Within each panel, rows correspond to values of $\alpha$ and columns to values of $\beta$ for the indicated values of echelon and $\omega$.

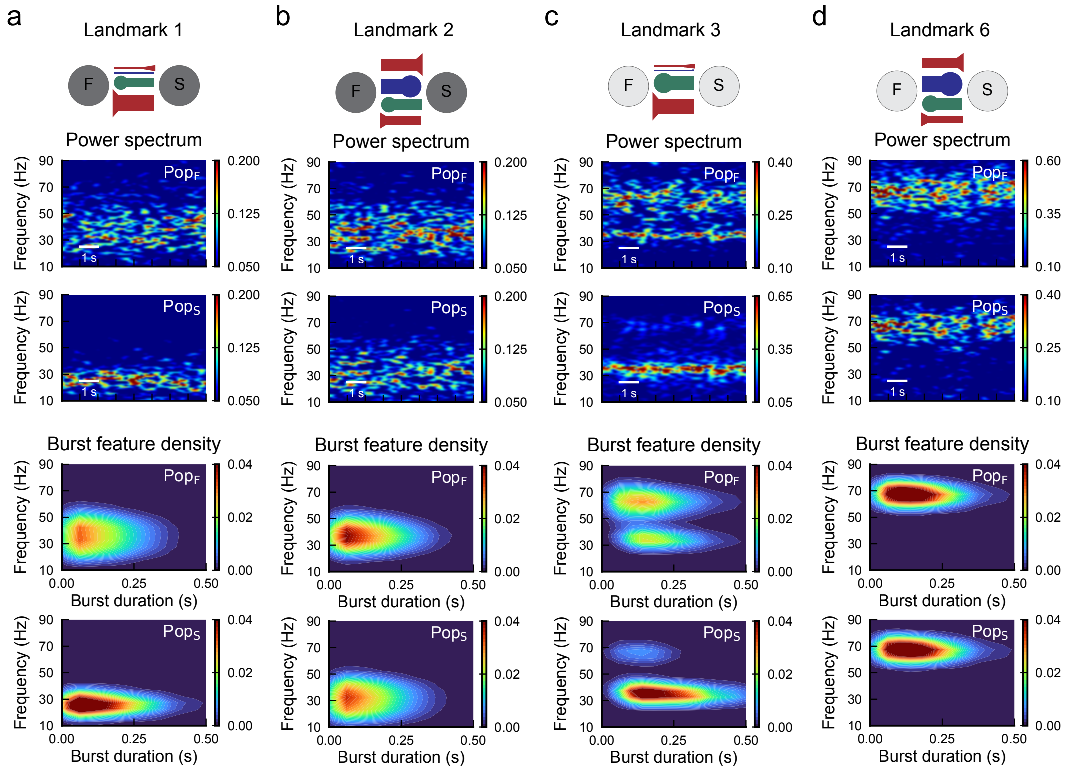


**Figure S5: Additional landmark configurations illustrate the diversity of coupled multi-frequency burst regimes.** Top, schematic of representative landmark circuits. Red arrows indicate excitatory projections, blue inhibitory projections from $\texttt{Pop}_F$ to $\texttt{Pop}_S$, and green inhibitory projections from $\texttt{Pop}_S$ to $\texttt{Pop}_F$. Arrow width scales with coupling strength, and node shading indicates echelon. Second and third rows, representative spectrograms for the $\texttt{Pop}_F$ and $\texttt{Pop}_S$. Fourth and fifth rows, joint burst-duration/frequency densities for each population. **a**, Landmark 1 shows broad-band transient bursting, with stronger fast components in the $\texttt{Pop}_F$. **b**, Landmark 2 resembles landmark 1 but with an upward shift of bursting frequency in the $\texttt{Pop}_S$. **c**, Landmark 3 shows mixed slow and fast bursting in both populations, with longer-lasting slow bursts. **d**, Landmark 6 is dominated by fast bursting in both populations.

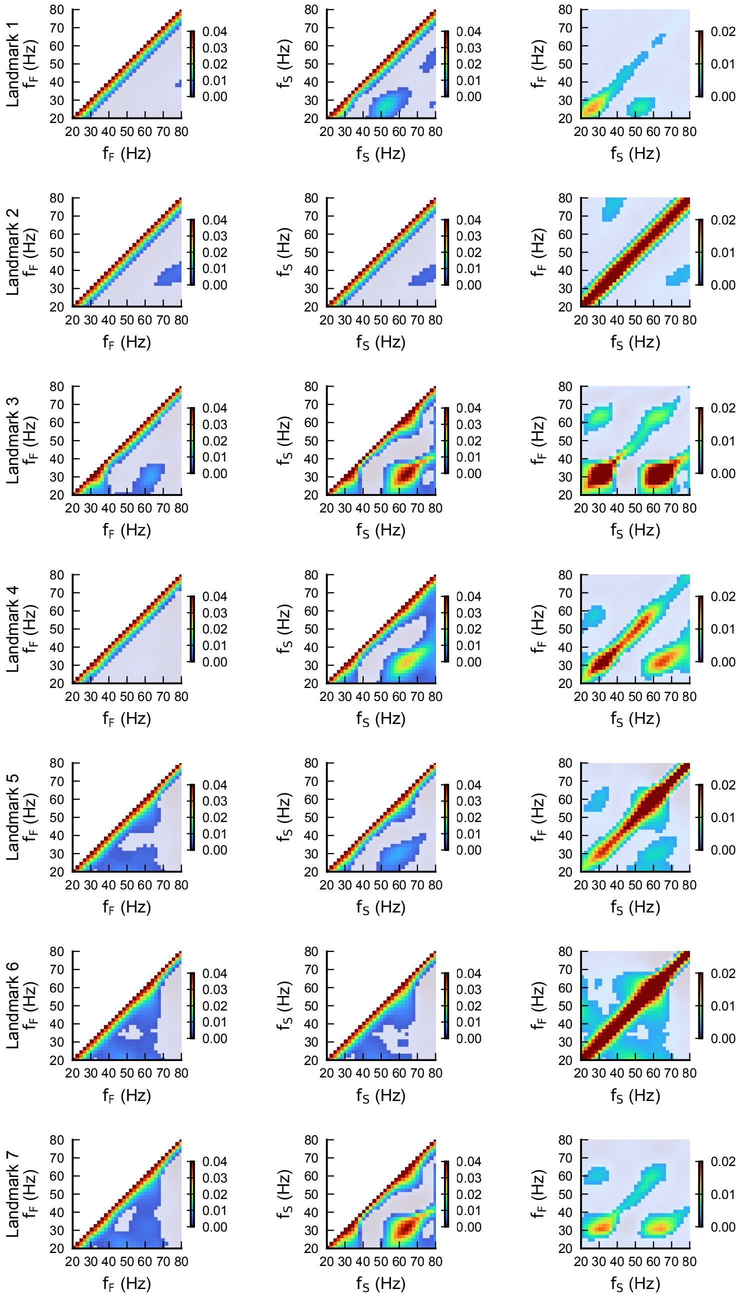


**Figure S6: Burst co-occurrence maps reveal structured coordination across landmarks.** For each landmark, the first and second columns show burst co-occurrence probabilities within $\texttt{Pop}_F$ and $\texttt{Pop}_S$, respectively, and the third column shows co-occurrence probabilities between the two populations. Frequencies are denoted by subscripts $F$ and $S$ for the $\texttt{Pop}_F$ and $\texttt{Pop}_S$. Co-occurrence was assessed from quantized spectrograms as in Fig. **??**. Opaque regions indicate frequency pairs whose co-bursting probability exceeded surrogate expectations, whereas non-significant entries are shown transparently. Across landmarks, burst coordination is structured and recurrent both within and between populations.

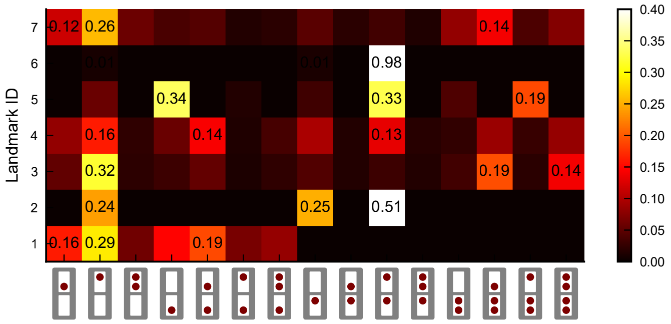


**Figure S7: The most frequent Multi-Frequency Oscillatory Patterns (MFOPs) differ across landmarks.** For each landmark, the three MFOPs with the highest occurrence rates are shown together with their corresponding probabilities of occurrence. Along the x-axis, each pictogram represents one such mask: the upper and lower rectangles correspond to $\texttt{Pop}_F$ and $\texttt{Pop}_S$, respectively and red dots indicate the presence of fast or slow burst components within each population (upper and lower positions inside each rectangle, respectively), as introduced in Fig. **??**a. The distribution of the most frequent MFOPs varies across landmarks, showing that each dynamical regime favors a distinct subset of coordinated multi-frequency burst configurations. Some masks are absent because the corresponding burst combinations are not expressed in a given landmark, or are not observed at all in the simulated dynamics.

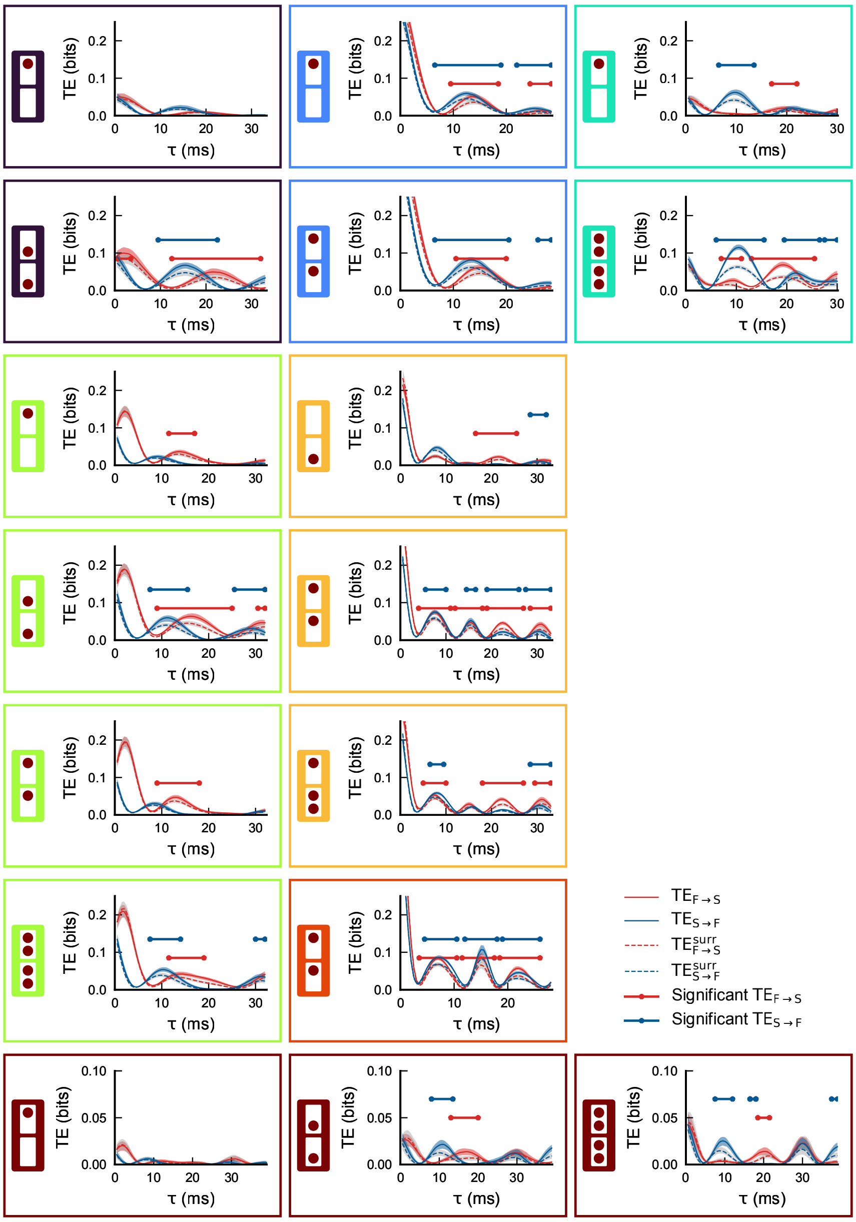


**Figure S8: Full lagged TE profiles underlying the Information Routing Patterns shown in the main analysis.** Each panel is framed by the color of the corresponding landmark, with the associated MFOP pictogram shown on the left. Red curves denote transfer entropy from $\texttt{Pop}_F$ to $\texttt{Pop}_S$ ($TE_{F \to S}$), and blue curves denote transfer entropy from the $\texttt{Pop}_S$ to the $\texttt{Pop}_F$ ($TE_{S \to F}$); dashed curves indicate the corresponding surrogate estimates. Shaded regions show confidence intervals. Horizontal red and blue segments indicate delay ranges in which TE is significantly greater than surrogate in the corresponding direction. These significant intervals are the basis of the IRPs summarized in Fig. **??**e.

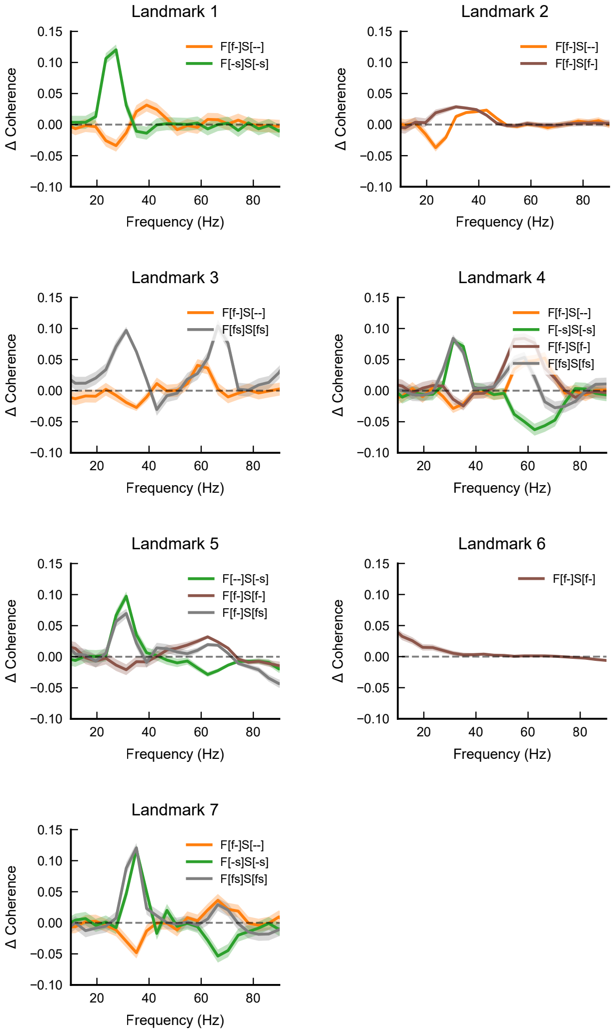


**Figure S9: MFOP-specific changes in coherence.** For each landmark, curves show the change in inter-population coherence relative to periods in which no MFOP is active. Uppercase $F$ and $S$ denote the $\texttt{Pop}_F$ and $\texttt{Pop}_S$, respectively, whereas lowercase $f$ and $s$ denote fast and slow burst components within each population; "-" indicates the absence of the corresponding component. Across MFOPs, coherence is selectively enhanced at frequencies represented in the corresponding burst mask, with the magnitude and spectral specificity of the effect varying across landmarks.

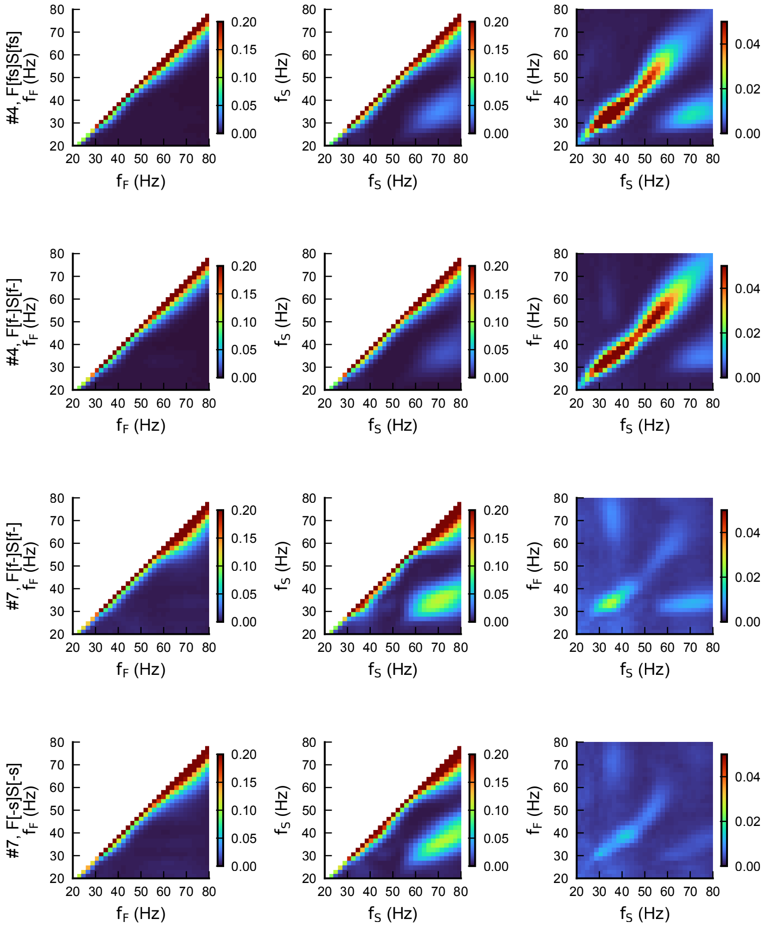


**Figure S10: MFOP-specific amplitude-amplitude coupling across frequencies.** Amplitude-amplitude coupling is shown for four example MFOPs across frequency pairs within the `Pop`$_F$ (left), within the `Pop`$_S$ (middle), and between `Pop`$_F$ and `Pop`$_S$ (right). Labels beginning with "#" indicate the landmark identity and associated burst mask. Amplitude was computed from the envelope of the Hilbert-transformed band-passed signals, and coupling strength was quantified by the mutual information between amplitudes at each frequency pair (see Methods). Related MFOPs show similar coupling architectures, with differences primarily in strength and spectral emphasis.

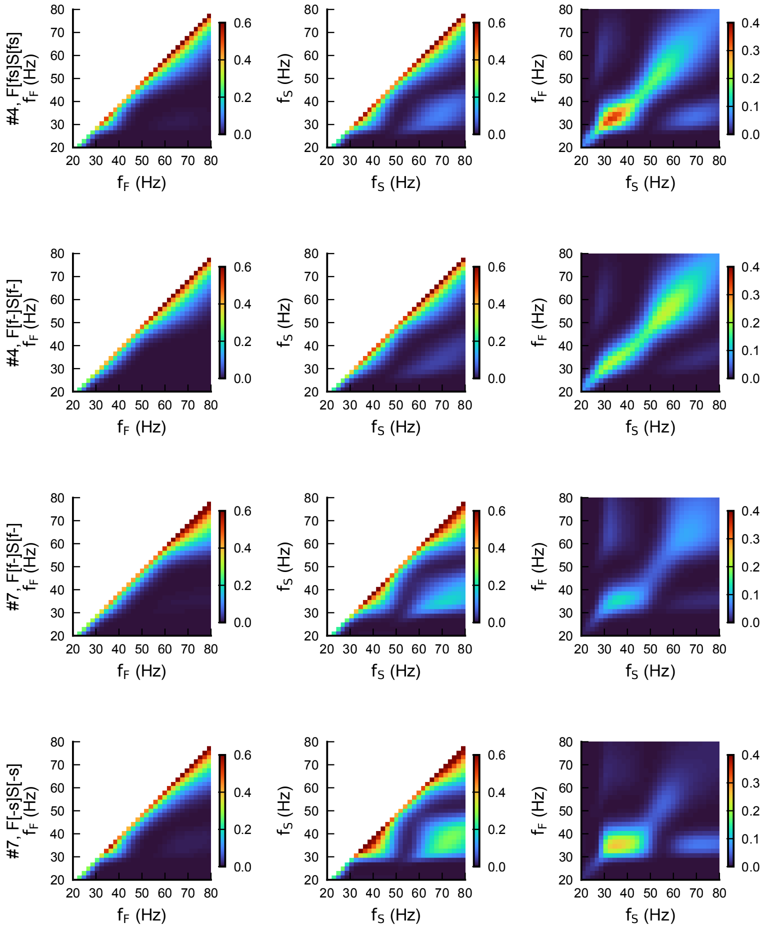


**Figure S11: MFOP-specific phase-phase coupling across frequencies.** Phase-phase coupling is shown for four example MFOPs across frequency pairs within the $\texttt{Pop}_F$ (left), within the $\texttt{Pop}_S$ (middle), and between $\texttt{Pop}_F$ and $\texttt{Pop}_S$ (right). Labels beginning with "#" indicate the landmark identity and associated burst mask. Phase was computed from the angle of the Hilbert-transformed band-passed signals, and coupling strength was quantified by the mutual information between phases at each frequency pair. MFOPs associated with different IRPs can nevertheless share related phase-coupling structures, consistent with a common landmark-dependent scaffold modulated by state.

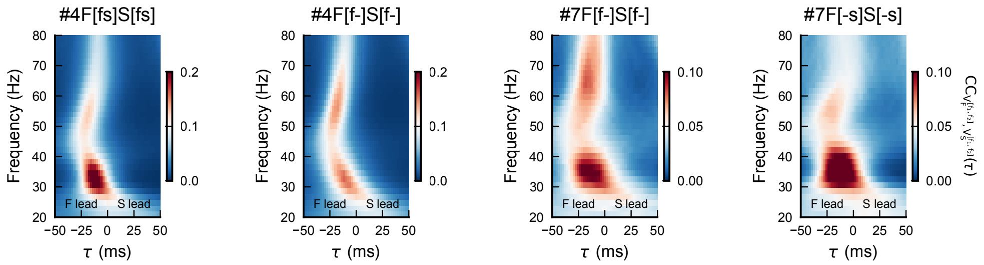


**Figure S12: MFOP-specific lagged cross-correlation patterns across frequencies.** Lagged cross-correlations between the LFP-like activities in $\texttt{Pop}_F$ and $\texttt{Pop}_S$ are shown for four example MFOPs as functions of time lag $\tau$ and frequency. Labels beginning with "#" indicate the landmark identity and associated burst mask. Negative $\tau$ indicates that the $\texttt{Pop}_F$ leads the $\texttt{Pop}_S$, whereas positive $\tau$ indicates the converse. MFOPs within the same landmark can exhibit distinct lag-frequency precedence structures, and similar burst masks can be associated with different precedence patterns in different landmarks.

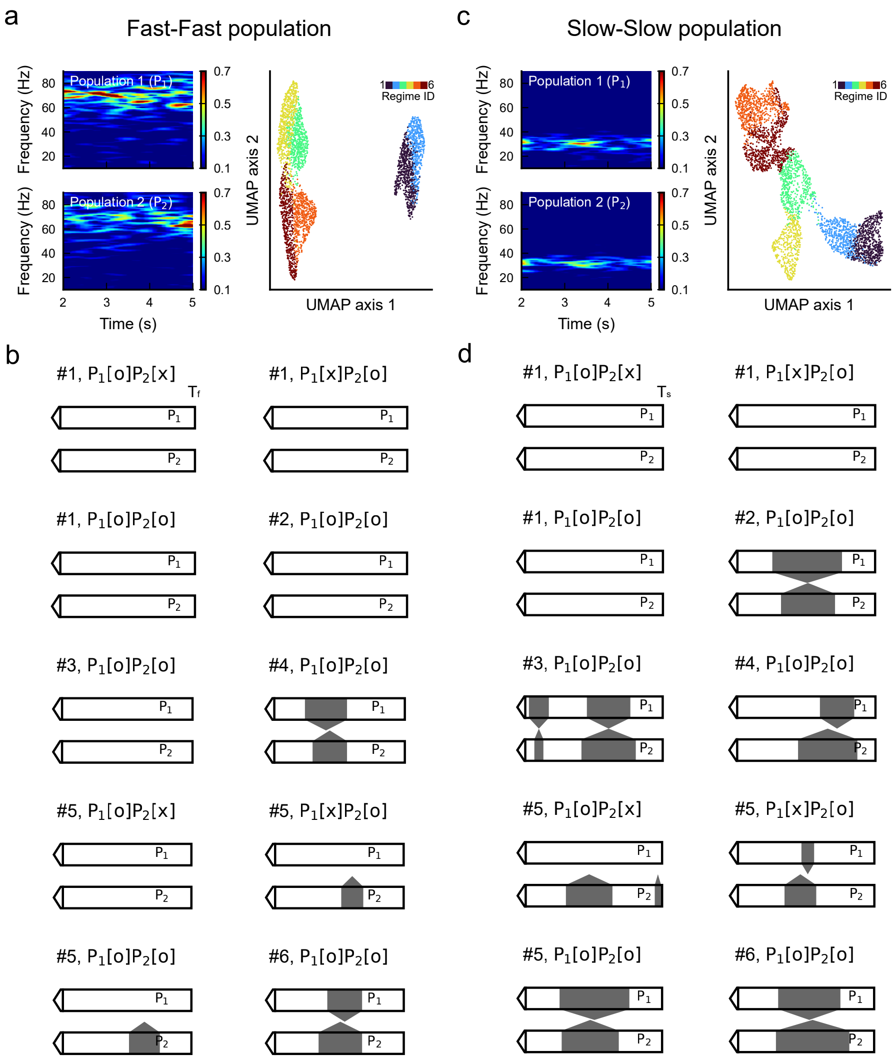


**Figure S13: Single-frequency coupled systems exhibit reduced MFOP and IRP diversity. a**, Example spectrograms for two coupled $\texttt{Pop}_F$ (left) and corresponding UMAP embedding of dynamical regimes (right). Structural parameters and analysis settings were matched to the main analysis, except for the symmetry constraints imposed by identical populations. **b**, IRPs identified for MFOPs in $\texttt{Pop}_F$-$\texttt{Pop}_F$ configuration. $P_1$ and $P_2$ denote the two populations. Because only one oscillatory timescale is available, MFOPs are defined solely by the presence or absence of fast bursts in each population, and the resulting IRP repertoire is limited. **c**, Same as in **a** for two coupled $\texttt{Pop}_S$. **d**, IRPs identified for MFOPs in $\texttt{Pop}_S$-$\texttt{Pop}_S$ configuration. As in $\texttt{Pop}_F$-$\texttt{Pop}_F$ case, restricting the system to a single oscillatory timescale markedly reduces the diversity of MFOPs and routing motifs relative to the main multi-frequency system.

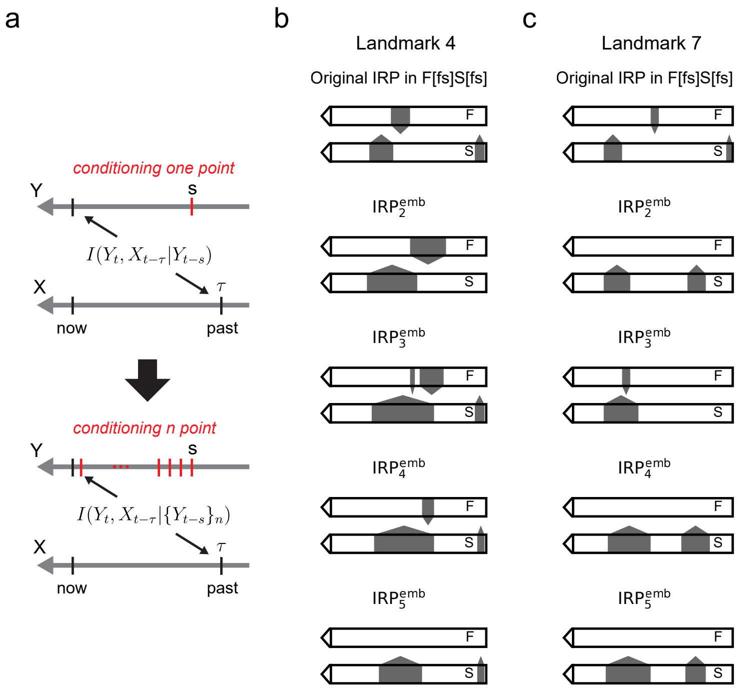


**Figure S14: IRPs are robust to richer embeddings of the target past in TE estimation. a**, Schematic illustration of TE estimation with increasing embedding dimension. For embedding dimension $n$, TE conditions the present of the target on an $n$-dimensional embedding of its past rather than on a single past time point (see Methods). When $n = 1$, this reduces to the TE definition used in the main analysis. **b,c**, IRPs obtained for two example MFOPs across embedding dimensions. Although arrow number and width may vary with embedding choice, the main routing delays and directions remain qualitatively stable, indicating that IRP structure is robust to the use of richer target-past embeddings.

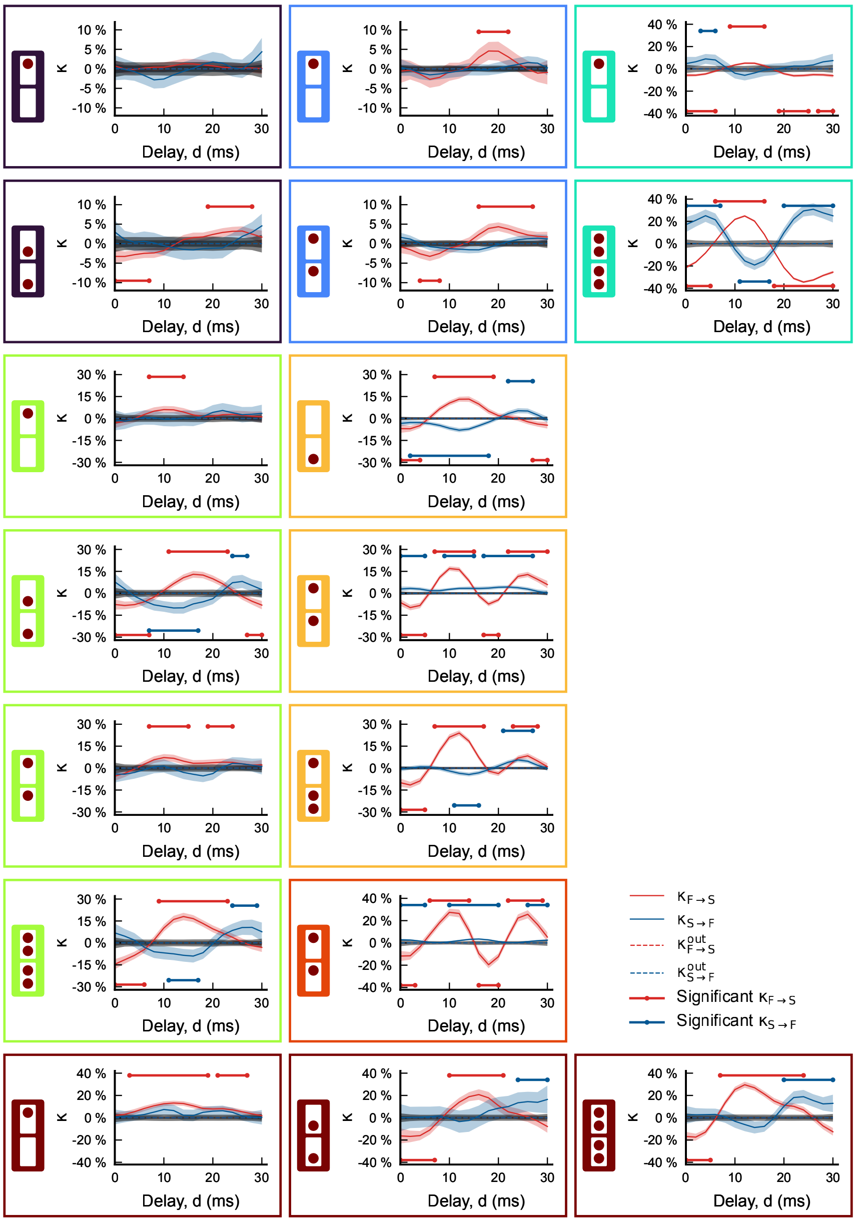


**Figure S15: Delay-dependent spike-transmission modulation curves for all states shown in the main analysis.** Each panel is framed by the color of the corresponding landmark, with the associated burst-mask pictogram shown on the left. Red curves denote $\kappa_{F\to S}$ and blue curves $\kappa_{S\to F}$; dashed curves indicate the corresponding out-of-state baselines. Shaded regions show confidence intervals. Horizontal red and blue segments indicate delay ranges in which transmission is significantly enhanced or suppressed relative to baseline in the corresponding direction. These significant windows are the basis of the spike-transmission barcodes summarized in Fig. ??f.

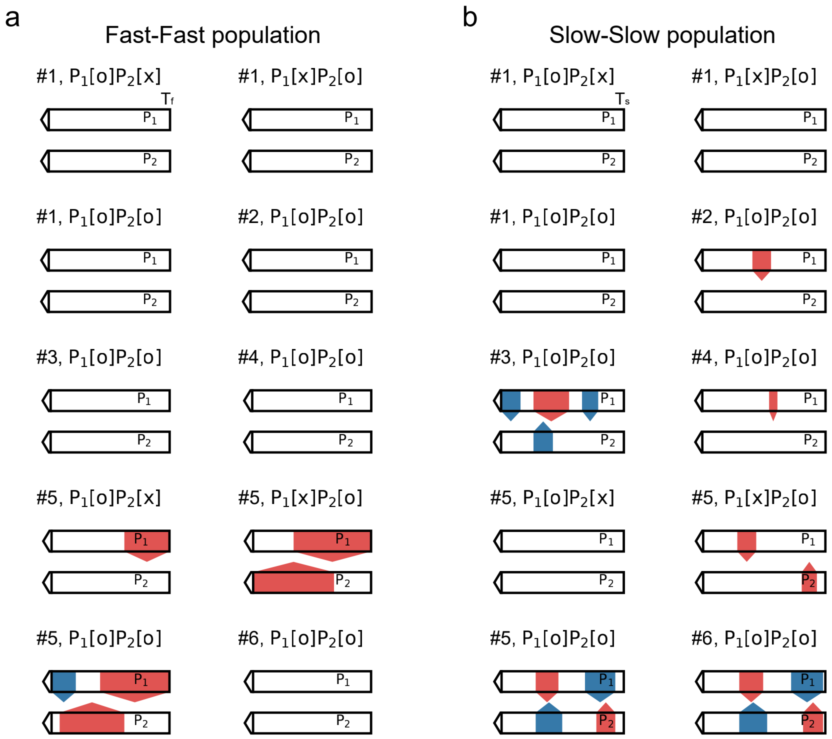


**Figure S16: Single-frequency coupled systems exhibit reduced diversity of spike-transmission barcodes. a**, Spike-transmission barcodes for coupled $\texttt{Pop}_F$-$\texttt{Pop}_F$ populations. The structures and states correspond to those shown in Fig. S13. Because only one oscillatory timescale is available, most states show little or no selective delay dependence, and the resulting barcode repertoire is limited. The end of each diagram corresponds to the fast oscillation period $T_f$. **b**, Same as in a for coupled $\texttt{Pop}_S$-$\texttt{Pop}_S$. Restricting the dynamics to a single oscillatory timescale markedly reduces the diversity of delay-dependent spike-transmission motifs. The end of each diagram corresponds to the slow oscillation period $T_s$.

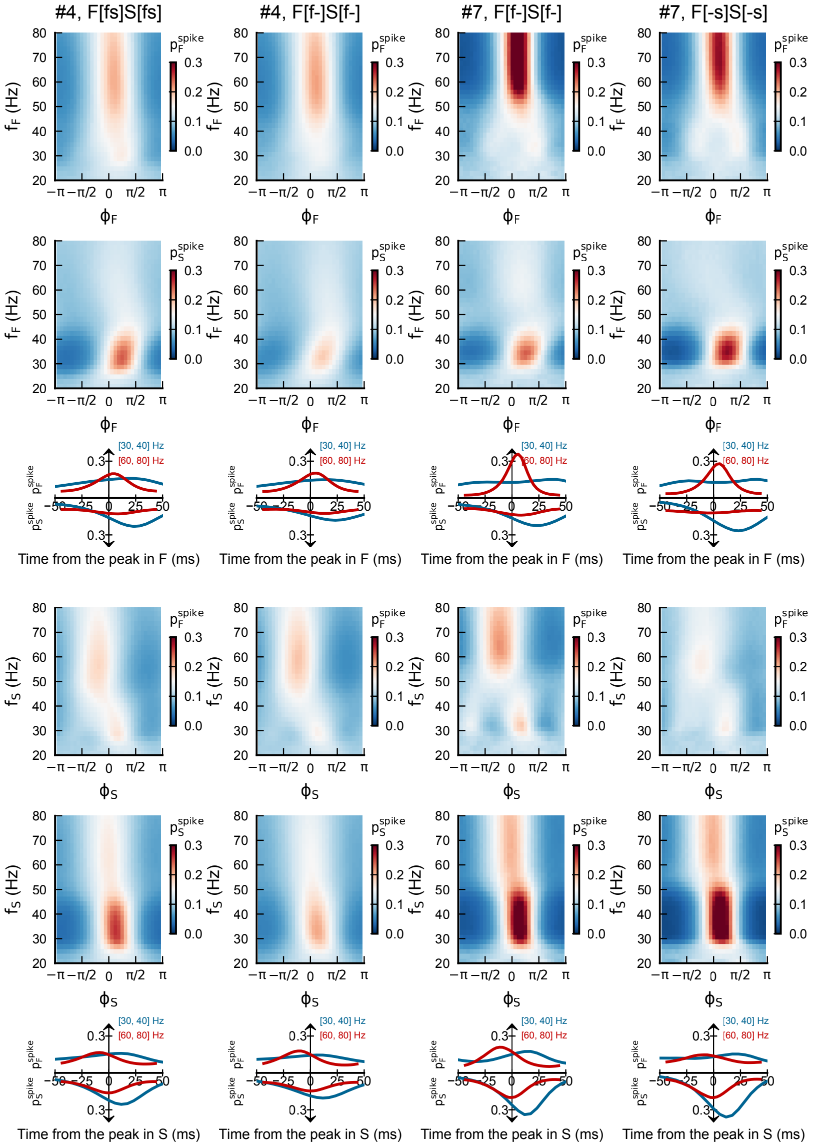


**Figure S17: Coordinated bursting states modulate background spiking probability as a function of oscillatory phase and frequency.** Columns correspond to four example coordinated bursting states, with labels indicating landmark identity and burst mask. Subscripts $F$ and $S$ denote quantities measured in $\texttt{Pop}_F$ and $\texttt{Pop}_S$, respectively. Rows 1–3 show spike probability ($p^{spike}$) in $\texttt{Pop}_F$ and $\texttt{Pop}_S$ as a function of the phase and frequency of oscillations measured in $\texttt{Pop}_F$; rows 4–6 show the same analysis with phase and frequency measured in $\texttt{Pop}_S$. The line plots convert phase to time from the oscillatory peak for the slow-frequency (blue) and fast-frequency (red) components. Coordinated burst states therefore impose structured phase-dependent modulation of population excitability, providing a dynamical background onto which delayed spike transmission is superimposed, with nonlinear additive effects.

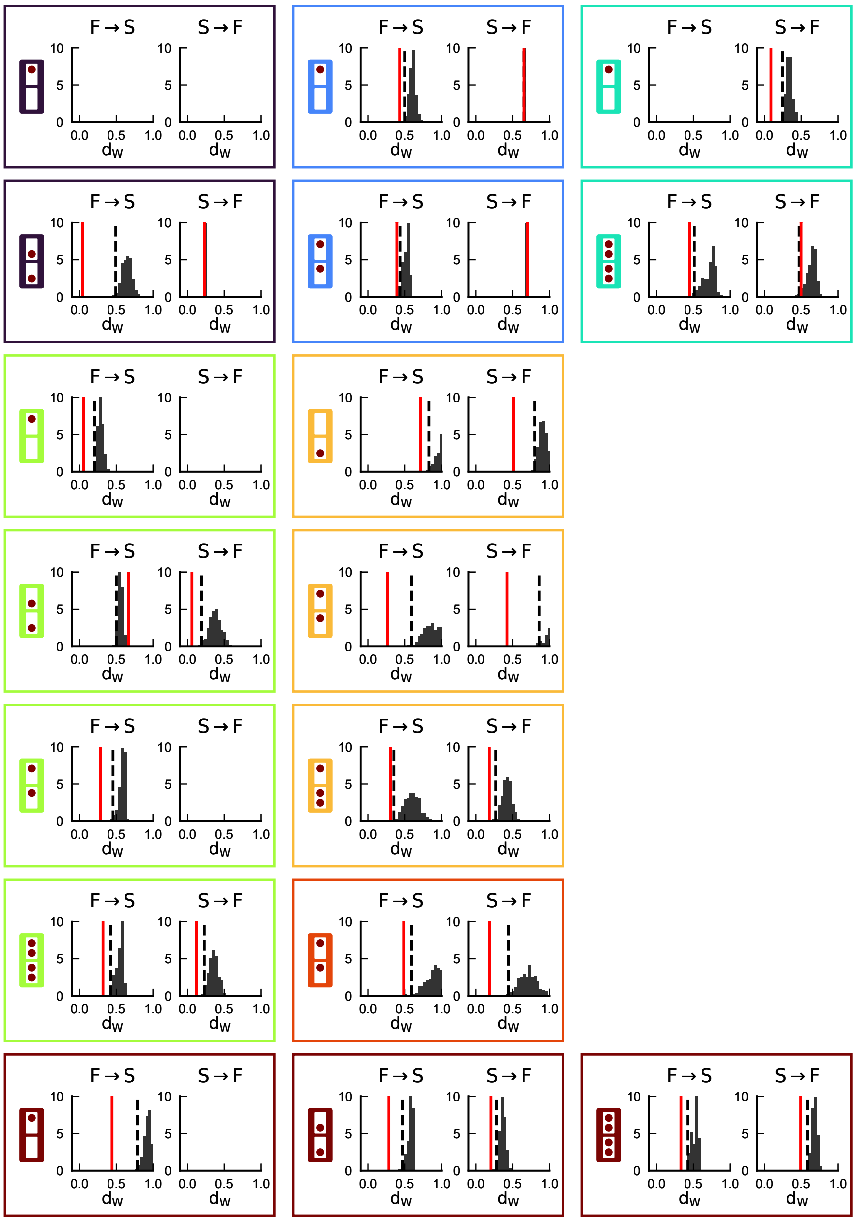

**Figure S18: Spike-transmission barcodes are more similar to matching TE-based routing motifs than expected by chance.** For each state shown in the main analysis, histograms display the surrogate distribution of Wasserstein distances $d_W$ between the TE-based IRP and shuffled spike-transmission barcodes preserving arrow number and width but randomizing their positions. Red lines indicate the observed distance for the matching state. Black dashed lines denote the significance threshold at $p = 0.01$. In most cases, the observed distance is smaller than expected from surrogate barcodes, indicating a statistically significant correspondence between mesoscale routing motifs and microscopic transmission patterns.
